# Hidden Interactions: Frequency-Dependence Emulates Selection-Driven Dynamics in Evolving Populations

**DOI:** 10.1101/2023.03.15.532871

**Authors:** Rowan J. Barker-Clarke, Jonathan Asher Pachter, Jason M. Gray, Sydney Leither, Maximilian A. R. Strobl, Jeff Maltas, Dagim Shiferaw Tadele, Michael Hinczewski, Jacob G. Scott

## Abstract

The temporal evolution of mutating pathogens in disease contexts arises from both the intrinsic properties of each subpopulation and the interactions among them, yet experimental inference often neglects the latter. Drug development studies commonly estimate selective advantages by comparing growth rates in monoculture, and in vitro monoculture dose–response curves are frequently used to justify or halt further investigation of a drug. Although many ecological models distinguish intrinsic from interaction-dependent growth rates, we show that simpler evolutionary game theory (EGT) frameworks can also be used to disentangle these contributions. We present a game-theoretic reparameterization of the replicator equation payoff matrix that separates intrinsic effects from interaction-driven contributions to frequency-dependent fitness. We also introduce an interaction–selection plot that facilitates the interpretation of the relative importance of between-population interactions compared with intrinsic evolutionary trade-offs. Using this framework, we map how interactions can mask, mirror, maintain, or mimic frequency-independent selection. We derive analytical conditions for these behaviors in both deterministic (replicator equation) and stochastic (Fokker–Planck–Kolmogorov) models, showing that simple conditions persist when mutation and noise are introduced. We validate these predictions using Wright–Fisher simulations. Applying our framework to published microbial and cancer co-culture data, we find that real systems span regimes dominated by either autonomous selection or interaction-driven effects, with interactions sometimes reversing or neutralizing frequency-independent fitness differences. Together, our results show that frequency-dependent effects can shape evolutionary dynamics in subtle and non-obvious ways, highlighting the importance of accounting for interactions when inferring fitness and predicting evolutionary outcomes.

## Introduction

Evolution – the changes in the frequency of alleles, or subpopulations – is often described through the lens of intrinsic monoculture fitness, which is the ability of a population with a single genotype to survive and reproduce in a given environment. [1] This idea underpins the original concept of the fitness landscape, a driving framework for understanding intrinsic fitness, selection, and evolution. [2–6] Within the peaks and valleys of the fitness landscape, the trajectory of evolution for a population is driven by mutational supply and the relative intrinsic fitness of accessible genotypes. [7–10] Frequency dependence (independent of mechanism) arises when the fitness of a genotype varies with its relative abundance in a mixed population. Frequency-dependent and ecological interactions within complex biological communities are now widely recognized to shape evolutionary outcomes, from microbial populations to cancer cell communities. [11–21] For example, Kaznatcheev *et al*. [21] demonstrated how alectinib-resistant and parental non-small cell lung cancer cells have different fitnesses across relative population frequencies and how the presence of fibroblasts or changing treatment results in distinct frequency-dependent functional changes in fitness (often termed evolutionary games). Subsequently, the interplay between genotypic fitness tradeoffs and frequency-dependent selection is of particular interest.

Although ecology does not traditionally model mutation, several ecological frameworks (e.g., Lotka-Volterra) do make clear distinctions between intrinsic (isolated) and frequency-dependent contributions to population growth. [22, 23] In contrast, evolutionary game theory (EGT) formalisms have thus far struggled to mesh with the monoculture-focused study of evolutionary tradeoffs. This difficulty arises in part because, although EGT payoff matrices can be derived straightforwardly from empirical growth-rate measurements, their origins in macroscopic biology, strategy, and behavior have limited direct connections to population-genetic concepts such as selection coefficients and monoculture fitness. Moreover, the common transformation to “game space” in the EGT literature emphasizes the outcomes of interacting strategies rather than the underlying structure and interdependencies of the payoffs themselves. We focus on how interaction and autonomous selection can be studied within the replicator equation and other density-independent models in frequency space. [24, 25]

In reframing the EGT payoff matrix in the language of selection, we encode a biologically intuitive relationship, while also incorporating frequency-dependent fitness interactions. Formalizing this space is of particular interest in the treatment of evolving, multicellular diseases such as cancer and complex infections. For example, while the heterogeneity of genotypes within a patient’s disease is accepted as a driver of both pre-existing and *de novo* resistance evolution, the presence of frequency-independent selection advantages and cell-cell interactions in mixed populations may both contribute to altering the efficacy of drugs. [16, 26] It is often the case that personalized treatment in evolving diseases eventually encounters resistance, the reasons for which likely include a combination of stochastic and interaction effects alongside both environmental and genotypic factors. [16, 27, 28]

In this manuscript, we begin with the deterministic replicator equation (RE). The replicator equation describes the rate of change of subpopulation *frequencies*, and while it can be derived from equations describing net population growth (see Supplementary Information), it does not describe population size or density-dependence. The RE and replicator-mutator equation (RME) both hold under assumptions of fixed-size or of exponential growth/decay. [24] These assumptions follow from the derivation of the replicator equation. Importantly, the rate of change of frequency in the RE model can be frequency-dependent, conceptualized using a game-theoretical payoff matrix, as we will employ here. While alternate models incorporate ecological variables, we focus on a modeling framework independent of population density. A key novelty in our work is the conceptualization of monoculture fitness as equivalent to the terms *a*_*ii*_ of the replicator equation payoff matrix. In contrast to the Lotka-Volterra model, a base monoculture growth rate in the replicator framework does not have to be assumed to be independent of the interactions within a sub-population.

One way in which interactions can be observed in co-culture experiments is to find the growth rate of each population at a range of starting fractions (**Fig 1A**). Changes in growth rates due to interaction effects are observed in a dose-dependent way within co-culture and can result in altered evolutionary outcome (**Fig 1B**). We propose four plausible regimes, defined by how frequency-dependent payoffs alter evolutionary outcomes relative to expectations based solely on intrinsic fitness differences (**Fig 1C**). Consider two genotypes in each plot, where the red has a higher *intrinsic* fitness than the blue. The first scenario, which we call *maintenance*, occurs when the interactions do not alter the outcome of evolution: a co-cultured mixture of two types will behave exactly as expected from monoculture, the red eventually outcompeting blue. This could be due to the absence of interactions, but as we will show below, there is also a set of non-zero interaction strengths that lead to maintenance. The second scenario is *masking*: two genotypes behave as if equally fit (neutral) when co-cultured, despite monoculture fitness differences. The third scenario is *mirroring*: where one is expected to dominate based on monoculture fitness (i.e., red versus blue), but the monoculture expectations are inverted in co-culture (blue outcompetes red to the same degree). The final scenario is *mimicry*: in which two genotypes have equal monoculture fitness but effectively behave as if one has a selective advantage when co-cultured. We focus on these four scenarios because they present interesting examples that clearly illustrate the potential confounding effects of frequency-dependence; interactions can be present even if a net effect is absent, and switching between monoculture and co-culture can either exacerbate or mitigate selective differences.

**Figure 1.**
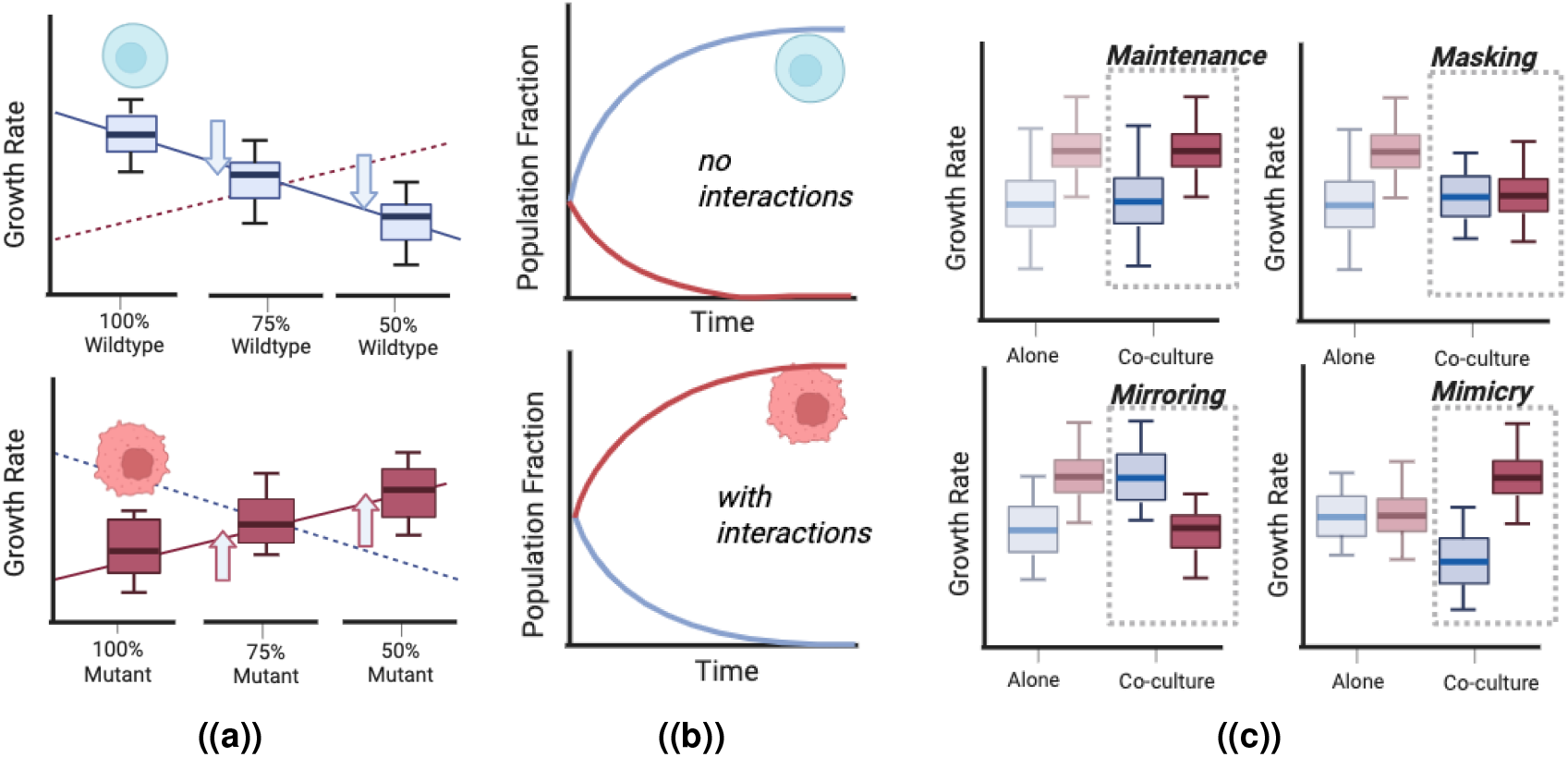
Intrinsic fitness differences combined with frequency-dependent interactions can result in counter-intuitive outcomes. Given the monoculture growth rates of two cell types, the null hypothesis is that the same rank and relative growth rate would be sustained. Interactions can be considered to be the deviation from this. Intrinsic fitness differences can be observed by comparing the growth rates of two different populations in monoculture. If the mutant has a higher growth rate than the wildtype in coculture, the mutant outcompetes the wildtype when growing in coculture. **(A)** It is common for experimental scientists to detect frequency-dependent interactions by measuring the change in the growth rates of populations in 50:50 co-culture compared to their monoculture growth rates. Frequency dependence can be seen explicitly when growth rates change depending on the ratio between the first and second types. **(B)** For the example in **(A)** where the mutant has a lower intrinsic fitness than the wildtype, without interactions, the wildtype would outcompete the mutant **(top). (C)** Given the importance of intrinsic fitness differences in evolutionary biology, we propose that specific classes can modify the resultant fitnesses in co-culture to result in different evolutionary dynamics and different steady states. We define four representative ways in which interactions confound monoculture expectations. Maintenance, where the original rank and relative distance are preserved. Masking, where original fitness differences are neutralised. Mirroring, where the intrinsic growth rate difference is reversed, and mimicry, where there is no intrinsic fitness difference, but one population outgrows another in co-culture.

Many papers have previously used game theory to model tumor growth and composition, with the presence of interactions between cells, including competing tumor and stromal cells, and the production of growth factors as a strategy. [29–35] Theoretical studies have also shown the potential of frequency-dependent selection to promote high mutation rates and accelerate evolution. [36, 37] Advancing upon prior formulations of frequency-dependent Wright-Fisher models [38, 39] and established co-evolutionary frameworks [40–42], we describe the four scenarios (**Fig 1C**) mathematically. Consistent with prior replicator equation models, we assume that cellular strategies are genetically encoded and inherited, such that payoffs in the evolutionary game represent a cell’s capacity to divide. Like these models, we assume that a cell’s strategy is determined by its genotype and inherited from its parent, such that the payoff in the evolutionary game reflects a cell’s ability to produce identical offspring. We also, for simplicity, have restricted our modeling approach to ignore density dependence. Remaining in frequency space also allows us to use the Fokker-Planck equation (also known as the Kolmogorov forward equation) to explore whether these behaviors exist (and can be defined) in the presence of mutation and genetic drift. [43, 44]

Motivated by increasing empirical evidence, but a lack of connection to fitness landscape and tradeoff studies, we focus on a game-theoretic framework that explicitly decomposes frequency-dependent fitness into cell-intrinsic and interaction contributions. This formalism enables the analysis of frequency dynamics of two co-cultured populations with varying intrinsic growth rates. Notably, the majority of assays in experimental evolution still measure and interpret the fitness of evolving populations primarily as an intrinsic property, independent of interactions between species.

Using both deterministic replicator and replicator-mutator equations, and the Fokker-Planck-Kolmogorov (FPK) description of evolving probability density in frequency space, we identify analytical conditions under which frequency-dependent interactions modulate the evolutionary dynamics to preserve, invert, amplify, or obscure the influence of intrinsic fitness differences. Our findings underscore the necessity of integrating frequency-dependent interactions into evolutionary models and evolutionary thinking, demonstrating that frequency-dependent selection can significantly reshape evolutionary trajectories. By analyzing 20 individual co-culture payoff matrices from seven published microbial and cancer co-culture experiments [21, 27, 45, 46], we observe both intrinsic-fitness-dominated and interaction-dominated regimes. We also show how interaction effects shift with environmental context, either reinforcing or counteracting intrinsic fitness differences. Our results highlight the importance of jointly considering frequency-dependent interactions and genetic fitness landscapes in evolution. These important findings have significant implications in interpreting experiments, designing treatments, and advancing work to control resistance evolution. For example, the application of control theory to steer populations across fitness landscapes must certainly account for the possibility that interactions influence the measured selection landscape. [47] Equally, omitting these factors could limit the success of approaches such as adaptive therapy, which involve mathematical optimization of tumor response with tailored on- and off-drug scheduling. [35, 48]

## Results

In the following, we will model the frequency dynamics of two populations: a wild-type population and a mutant population, using ideas from evolutionary game theory. We will study the dynamics and steady states of the *proportion* (*x*_*m*_ = *x*) of the mutant population (sometimes denoted the allele fraction). We will use the replicator equation formalism and the game-theoretical payoff matrix, which define frequency-dependent fitness and selection.

### Cell-intrinsic and frequency-dependent components can be logically separated within the payoff matrix

We ask, initially, how the traditional payoff matrix from game theory can be expressed in terms of the intrinsic selection coefficient? By decomposing the pay-off matrix into the contributions of cell intrinsic fitness and interaction effects, we can study the impact that each has on long-term evolutionary outcomes. To decompose the parameters describing frequency-dependent evolution in the game theory setting, we use the intuition that for there to be no frequency-dependence of the growth rates, each row of the payoff matrix must be constant, such that the payoff (growth rate) doesn’t depend on the populations present (**Box 1**).

Our decomposition of the payoff matrix (**Eq**. (B1)) is designed to explicitly separate the genotypic baseline growth rate from the type-interaction effect. In EGT, there is no additional isolated intrinsic growth rate term. We decouple an intrinsic term for each population that remains constant across the rows of the payoff matrix. This creates coupling within the EGT payoff matrix that is (a) intuitive and (b) has not been previously studied.

Consequently, when the interaction terms approach zero, (*α*_*wm*_, *α*_*mw*_ → 0), there are no between-population interactions and no frequency dependence. Additionally, when analyzing these dynamics, we have set the wild-type monoculture growth rate (*g*_*w*_) to 1. Should the explicit temporal scale be of interest to the reader, it is straightforward to reintroduce the scaling factor and recover this.

#### Box 1: Deterministic game theoretical framework and the payoff matrix

In the payoff matrix for two distinct cell populations undergoing replicator dynamics, the diagonal terms, *a*_11_ and *a*_22_ from **Eq**. (9), are understood as the reproductive fitnesses, and thus can be understood as the traditional monoculture growth rates of each population; that is, the reproduction rate of population 1 is experimentally measured in the presence of itself. The off-diagonal elements reflect the growth rate of the population in the presence of the other cell type. We model this, in our novel framing of the replicator equation (RE), as the population’s intrinsic growth rate modified by extrinsic cell-cell interactions. To illustrate this clearly, we represent and decompose the classical payoff matrix, *A*, to show the inter-dependence of coefficients as follows:

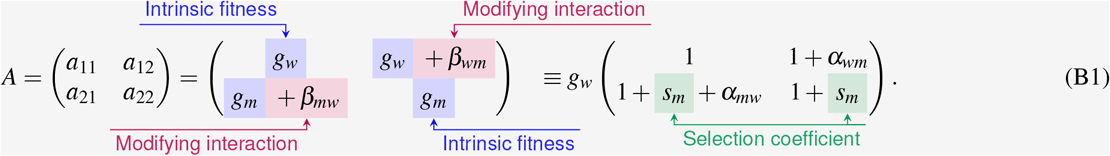

Here *g*_*w*_ and *g*_*m*_ are the per capita growth rates (effective fitness) of the wild-type and mutant cells, respectively, *β*_*wm*_ is the per capita fitness difference of the wild-type in the presence of the mutant population, and *β*_*mw*_ is the fitness difference of the mutant in the presence of the wild type population. In the second line, we have normalized all terms relative to the wild-type growth rate, defining the mutant selection coefficient *s*_*m*_ = *g*_*m*_/*g*_*w*_ − 1 and two interaction coefficients *α*_*wm*_ = *β*_*wm*_/*g*_*w*_, *α*_*mw*_ = *β*_*mw*_/*g*_*w*_.

After this variable transformation, the well-known replicator equation (**Eq**. (10)) for the mutant fraction *x* in our model can be rewritten as follows:

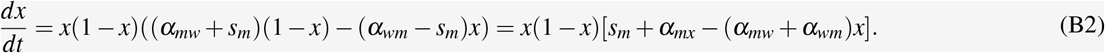

Using both the dynamics described by this equation and the possible steady-state solutions, we define the maintenance, mirroring, masking, and mimicry regimes in the text.

We then use this parameter transformation to rewrite the replicator equation. This equation describes the change in the mutant proportion (*x* ∈ [0, 1]), not absolute population size, over time. [49] In this parameterization, the dynamics and evolutionary outcomes (stationary state(s)) of competition between two populations can be written as functions of *α*_*wm*_, *α*_*mw*_, and *s*_*m*_. In this model, any specific evolutionary trajectory depends on the initial condition (the starting mutant fraction, *x*_0_), the stationary solutions and their stability, and the sign of 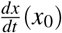.

To further visualize the relative contributions of interaction terms and intrinsic fitness differences, we introduce the “interaction-selection” plot (**Fig 2**), in which the axes are the ratios of extrinsic vs. intrinsic fitness of the wildtype and mutant type, respectively (*α*_*mw*_/*s*_*m*_, *α*_*mw*_/*s*_*m*_). In this representation, the unit circle is the region where intrinsic selection differences are dominant, whereas outside this region, between-population interactions are the dominant contribution to the fitness of the population. To understand how these components combine to influence the long-term dynamics, we overlay the long-term steady-state onto this plot (corresponding to the four quadrants of the traditional game-space plot). We can see that when interaction contributions are small (the ratio of interaction to selection is inside the unit circle), the mutant population wins due to its higher assumed intrinsic fitness. However, as we change the strength and direction of interactions, all four evolutionary scenarios are achievable, *despite the intrinsic fitness difference remaining unchanged*. The boundaries between the EGT quadrants (regions of parameter space with qualitatively similar equilibrium solutions) in the reparameterization occur along the lines *α*_*wm*_ = ™*s*_*m*_ and *α*_*mw*_ = *s*_*m*_. For example, the cooperation quadrant (top right) is defined by *α*_*wm*_ > *s*_*m*_ and *α*_*mw*_ > ™*s*_*m*_. In this way, we can see that as we vary the difference in intrinsic fitness (*s*_*m*_), the quadrants of different evolutionary outcomes are translated along the second diagonal and overlap exactly with the primary axes when *s*_*m*_ = 0. The interaction-selection plot also highlights how two systems can occupy the same location in the EGT game space (e.g., *G*_1_ and *G*_2_) with the same relative invasion fitness, but differ vastly in the underlying payoff contributions that brought them there.

**Figure 2.**
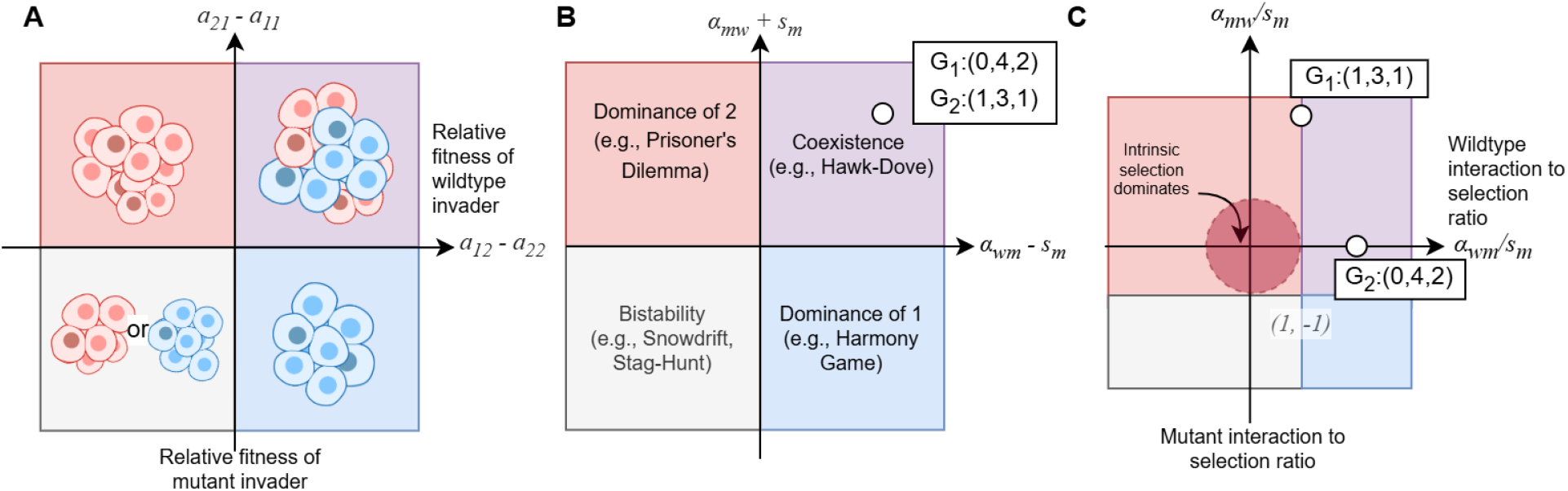
Resultant classes of evolutionary dynamics in the 2-player replicator game as a function of relative fitness and interaction strength. Illustrative diagram in which the color of the quadrants and position of the boundaries in these plots refer to four different universality classes (regions where the evolutionary stable state phase space is similar), as defined for deterministic evolutionary games. Boundaries between the classes in all phase spaces occur along the lines *α*_*wm*_ = ™*s*_*m*_ and *α*_*mw*_ = *s*_*m*_ and thus are mirrored accordingly in the interaction-selection plot depending on its sign. **(A)** Illustration of game dynamics between two populations (wild-type and mutant cells). Evolutionary game theory between two players results in four types of dynamics. The game space reflects distinct evolutionarily stable states corresponding to the four quadrants in relative fitness space (*a*_12_ − *a*_22_, *a*_21_ − *a*_11_). In the biological case, both the cell-intrinsic and between-population interaction influences on growth rate come together in co-culture to produce novel dynamics. The observed dynamics rely not only on the isolated behaviour of cells but also on the precise balance between intrinsic and extrinsic factors. In the top left quadrant, the wild type dominates. In the top right quadrant, a heterogeneous mixture is promoted. In the bottom right quadrant, the resistant mutant dominates. Lastly, in the bottom left quadrant, coexistence is unstable, and when the population is not exactly at the unstable fixed point (coexistence fraction), the population is driven to the nearest stable fixed point (all wild types or all mutants). **(B)** In the cell-centered decoupled payoff framework, the (*a*_12_ − *a*_22_, *a*_21_ − *a*_11_) axes and conditions on the game parameters for each quadrant are rewritten in terms of the intrinsic fitness difference (selection coefficient) and the interaction coefficients. *G*_1_ and *G*_2_ represent two payoff matrices encoded by (*α*_*ij*_, *α*_*ji*_, *s*_*m*_). These two systems share the same location in the invasion fitness plot. **(C)** In our interaction-selection plot, a system’s position is plotted as a function of interaction strengths (*α*_*ij*_) divided by the intrinsic selection coefficient *s*_*m*_. The coordinates in our mutant and wild-type system are (*α*_*mw*_/*s*_*m*_, *α*_*wm*_/*s*_*m*_). The previous *G*_1_ and *G*_2_ are no longer in the same position. The regions are labeled under the assumption that *s*_*m*_ > 0 and the mutant is fitter than the wild type. We highlight the unit circle in this space as the region where intrinsic selection is stronger than interaction. In this region, the fitter monoculture population (in this example, the fitter population is the mutant with *s*_*m*_ > 0) remains dominant.

Biologically, while intrinsic selection advantages can be present, cells within heterogeneous populations must survive selection amid interactions with other types. Taking into account the effects of both interaction and mutational forces, survival of – or competitive exclusion by – a sub-population may play out in a variety of ways: the maintenance of existing intrinsic selection advantages, the masking of selection differences, or the mimicry of intrinsic selection in its absence. Furthermore, one can also observe mirroring, the complete inversion of selection (direction and magnitude) due to interactions. These four interaction classes are of interest when interpreting the results of co-evolutionary dynamics in experiments.

### Game-modification of the intrinsic selection coefficient

Frequency-dependent dynamics – as described in evolutionary game theory – between wild-type and mutant alleles result in frequency-dependent selection. This is, by necessity, related to both the intrinsic population growth rates of cells (as measured in monoculture) and the interactions between cell populations *α*_*wm*_ and *α*_*mw*_. We hypothesized that the four interaction classes (as illustrated in **Fig 3**) are the result of specific relationships between the inter-species interaction-dependent contribution to selection and the intrinsic selection coefficient, which are in turn determined by the relationship between the interaction coefficients. To characterize this, one must ask what form the frequency-dependent selection coefficient would take?

**Figure 3.**
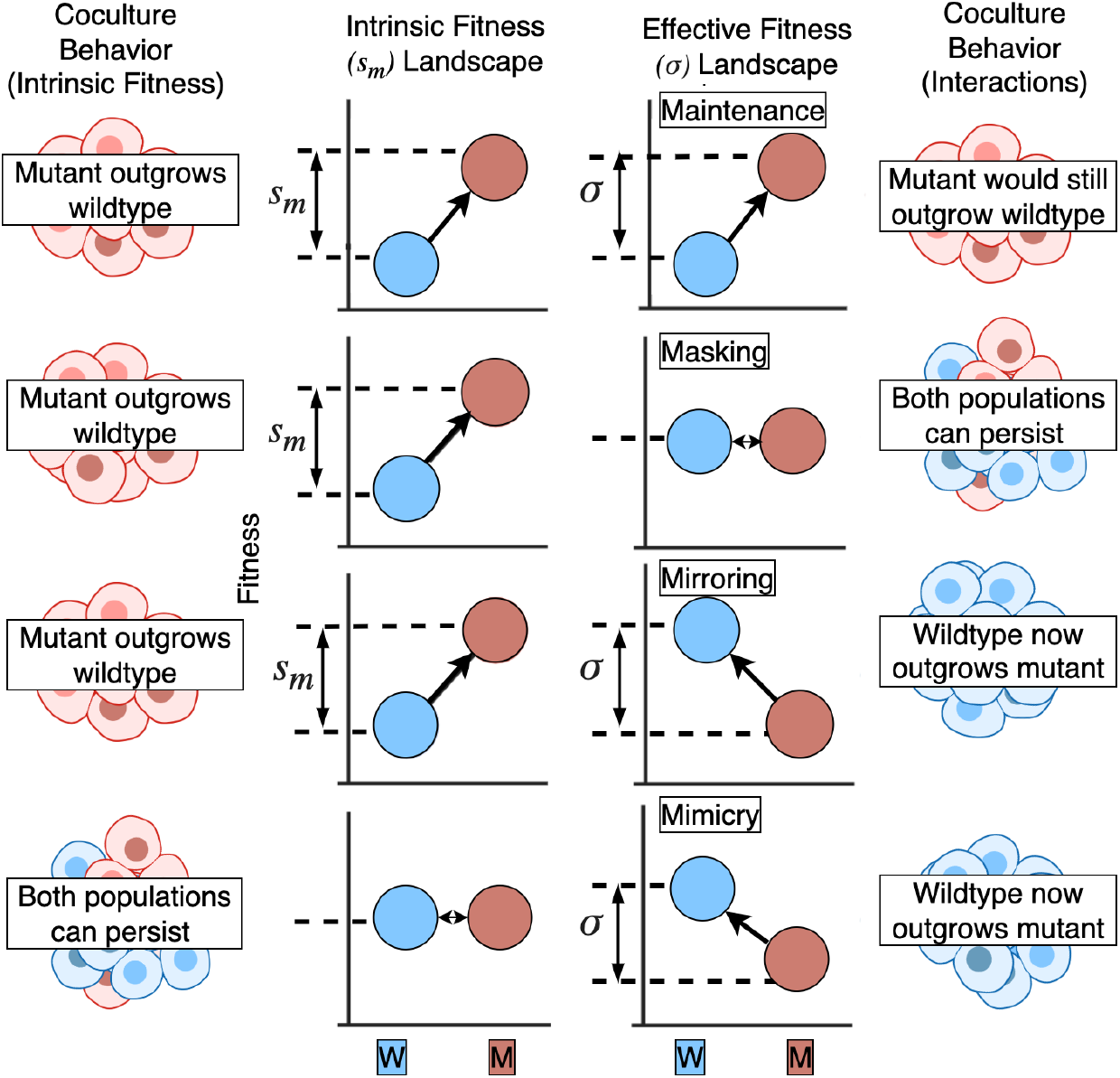
Effective fitness landscapes and definitions of maintenance, masking, mirroring, and mimicry in terms of the selection coefficient. Illustrative diagram of intrinsic monoculture fitness landscape representing the relative fitness difference (*s*_*m*_) of two different cell types (**left**), with two genotypes (wildtype and mutant) represented and an arrow from the lower to higher fitness genotype. Under this selection coefficient or fitness difference, there is a baseline expectation of how the evolutionary dynamics of a mixed population (blue and red) would play out: the proportion of the fittest genotype in the population increases towards 100%. For example, if the mutant (red) has a higher fitness in the effective fitness landscape, the population should be dominated by mutants. For co-cultured populations, interactions can modify the fitnesses and result in different evolutionary dynamics in co-culture. This effective fitness landscape has a new selection difference of *σ*. Thus, the evolutionary outcomes are modified by interactions, and the four rows show four representative ways in which interactions confound monoculture expectations: maintenance, masking, mirroring, and mimicry.

To discuss frequency-dependent dynamics relative to the intrinsic selection coefficient, we derive a new frequency-dependent effective selection coefficient using game-theoretical formalism. Note that, when the interaction terms go to zero (*α*_*wm*_, *α*_*mw*_ → 0) in **Eq**. (B2), the term in parentheses reduces to the selection coefficient *s*_*m*_. We thus define the frequency-dependent effective selection coefficient 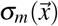 to be that term in parentheses:

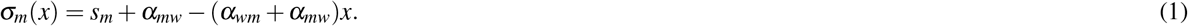

With this frequency-dependent effective selection coefficient defined, we can compare intrinsic (non-interacting) selection to frequency-dependent selection across varying intrinsic and interaction strengths.

### Systems with non-trivial interactions can reproduce frequency-independent dynamics

It is important to note, that when the selection coefficient is non-zero and there are no interaction terms (*α*_*wm*_, *α*_*mw*_ → 0), the replicator equation reduces, such that there are only two steady-state solutions, *x*_∞_ ∈ {0, 1}. This is frequency-independent deterministic evolution. When selection, *s*_*m*_, is non-zero, one population eventually outcompetes the other, the mutant proportion becoming *x* = 1 (100%) or *x* = 0 (0%) depending on the sign of *s*_*m*_.

In this replicator equation framework, we can define the concepts of maintenance, mirroring, masking, and mimicry via the preservation or inversion of the selection coefficient. In all of these cases, frequency dependence is removed despite the presence of non-zero interaction terms. As such, these solutions result in identical dynamics to the reference non-interacting case across time for all starting fractions. The alteration of the effective fitness landscapes through interactions, producing maintenance, masking, mimicry, and mirroring, are illustrated in **Fig 3**.

Maintenance occurs when the frequency-dependent selection coefficient is kept constant, *σ*_*m*_(*x*) = *s*_*m*_. In the second regime, fitness differences between a wild-type and a mutant in monoculture may be counteracted by interactions when they are grown in co-culture. We call this the *masking* scenario, where intrinsic fitness differences are neutralized by interactions. Masking occurs when the frequency-dependent selection coefficient, *σ*_*m*_(*x*) = 0. The reverse case is also possible, where wild types and mutants may have a neutral fitness relationship in monoculture, but when grown in co-culture, there is a selective fitness difference. Mimicry occurs when there is a zero intrinsic selection coefficient (*s*_*m*_ = 0), but a non-zero frequency-dependent selection coefficient, *σ*_*m*_(*x*) = *s*^′^. Mirroring occurs when the direction and magnitude of the selection coefficient are inverted during frequency-dependent selection, *σ*_*m*_(*x*) = −*s*_*m*_.

When these conditions upon the frequency-dependent *effective selection coefficient* occur, the resultant dynamics are indistinguishable from the respective frequency-independent cases. These regimes and the restrictions they place on the interaction coefficients are summarised as follows,

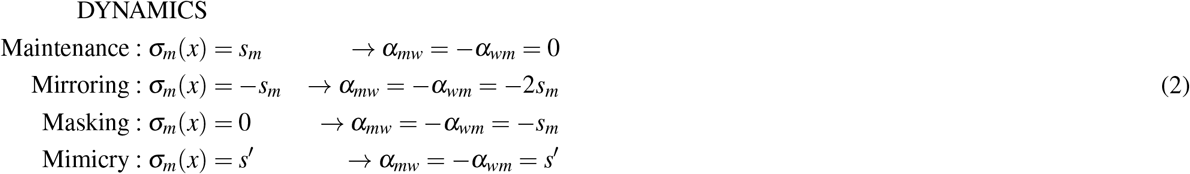

The full derivations of these conditions can be found in the Supplementary Information. Within the four cases defined, it is important to emphasize that the only possible *maintenance* is the trivial solution, *α*_*mw*_ = *α*_*wm*_ = 0. Across the three other non-trivial cases, the deterministic dynamics of the interacting system are indistinguishable from those of a non-interacting reference system, independent of starting conditions. In general, whenever *α*_*mw*_ = −*α*_*wm*_, the frequency-dependence is lost from **Eq**. (1), and the effective selection coefficient becomes *s*_*m*_ + *α*_*mw*_ Thus, when observing the dynamics of a system, it is possible for dynamics consistent with a non-interacting system of intrinsic selection coefficient *s*_*m*_ + *α*_*mw*_, to be produced by an interacting system with intrinsic selection coefficient *s*_*m*_ and interaction parameter *α*_*mw*_ = −*α*_*wm*_.

### Interacting systems can reproduce non-interacting equilibrium solutions

When we observe systems, for example, in experiment, or *in vivo*, we are not always able to observe the full *temporal dynamics* of a system, and effective selection is sometimes inferred from the steady state of a system. Having shown that we can define our interaction classes using the equivalence of the replicator equations, it is also possible to define these regimes via the equivalence of evolutionary outcomes (stationary solutions). We ask the question: for what (less strict) parameter ranges do deterministic co-evolutionary systems with interactions reproduce steady-state solutions that are indistinguishable from possible outcomes of systems without interactions?

A definition of *masking* in this case requires the production of the same long-term behavior as that obtained when *s*_*m*_ = *α*_*mw*_ = *α*_*wm*_ = 0, in which case the right-hand side of the replicator equation is identically zero, indicating that the mutant frequency remains for all time at its initial value. The conditions for masking in the long-term behavior of systems obeying the replicator equations turn out to be the exact same as the conditions for masking in the full temporal dynamics, given in **Eq**. (2).

As for maintenance, mirroring, and mimicry, note that the interactionless replicator equation has fixed points at *x* = 0 and *x* = 1, with the sign of *s*_*m*_ determining which is stable (*s*_*m*_ > 0 means the mutant will dominate and *x* = 1 is the stable long-term behavior, and vice versa). With interactions included, however, a third potential stationary frequency *x*^∗^ = (*s*_*m*_ + *α*_*mw*_)/(*α*_*mw*_ + *α*_*wm*_) emerges, in addition to the already extant values at 0 and 1. If this stationary frequency is physically accessible with 0 < *x*^∗^ < 1, the long-term behavior of the interacting system is fundamentally different from the interaction-free case, with either stable coexistence or bistability. If, however, *x*^∗^ ≥ 1 or *x*^∗^ ≤ 0, the long-term behavior will still be one stable accessible fixed point corresponding to either fixation or extinction of the mutant, since values of *x* less than 0 or greater than 1 are physically inaccessible. The condition that *x*^∗^ ≤ 0 or *x*^∗^ ≥ 1 can be expressed as:

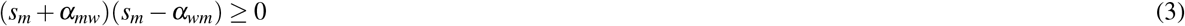

This condition must be satisfied for any of the four emulation categories to be possible, including masking (for which the equality is satisfied). Under this condition, the quantity *σ*_*m*_(*x* = 1/2) = *s*_*m*_ + (1/2)(*α*_*mw*_ − *α*_*wm*_) cannot be zero, and in fact determines the long-term stable fixed point: *σ*_*m*_(1/2) > 0 stabilizes *x* = 1, and *σ*_*m*_(1/2) < 0 stabilizes *x* = 0.

Combining these conditions, we find a broad range of interaction parameters for which maintenance, mirroring, masking, and mimicry can occur at steady state, summarised here (see Supplementary Information for full derivations). Let 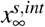 be the stable, accessible steady-state frequency with intrinsic selection coefficient *s* under interactions and let 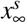 be the stable steady-state frequency with intrinsic selection coefficient *s* without interactions:

STEADY STATE

(*s*_*m*_ + *α*_*mw*_)(*s*_*m*_ − *α*_*wm*_) ≥ 0 satisfied in all cases, plus the following:

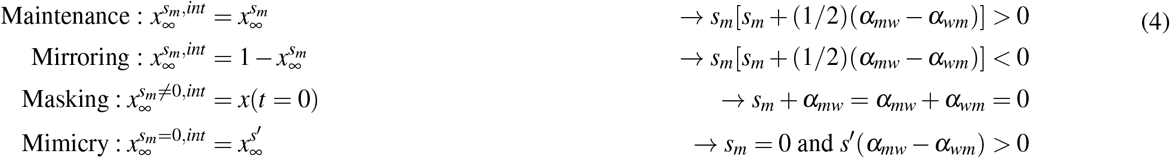

Thus, maintenance, mirroring, masking, and mimicry of the steady state of the deterministic replicator equation are possible, in which the long-term behavior of the interacting system is *indistinguishable* from the steady-state solution of the non-interacting system. For masking, the same equality constraints exist for full temporal dynamics emulation as for steady-state emulation. However, the parameter constraints for *maintenance, mirroring*, and *mimicry* in steady-state are inequalities – a broad range of interaction parameters can lead to emulation of non-interacting replicator systems when only studying the long-term behavior.

### Introducing mutation and genetic drift

The introduction of mutations transforms the deterministic replicator equation (RE) into the replicator-mutator equation (RME); in the Supplementary Information, we review the deterministic RME. In the main text, we will present a Fokker-Planck-Kolmogorov (FPK) equation for our system to incorporate both mutation and stochasticity into evolutionary dynamics (see **Box 2** for details). [43, 50] In the infinite population limit, the FPK equation reduces to a form of RME.

#### Box 2: Stochastic formalism

In a Fokker-Planck-Kolmogorov (FPK) description of our system, we denote *ρ*(*x,t*) as the probability density function of a population having mutant fraction *x* at time *t*. As shown by Kimura, [51] Wright-Fisher dynamics of a large (constant) population with mutation and selection of moderately weak magnitude can be described via the following diffusion equation,

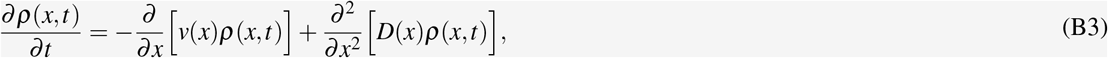

The first term on the right describes the deterministic effects of mutation and selection, which shift the peak of *ρ*(*x,t*) over time, while the second term leads to a diffusive broadening of the distribution due to demographic noise (sampling from a large but finite population of size *N* ≫ 1). While the diffusion term, *D*(*x*), remains constant (since it does not depend on selection), the original drift term *v*(*x*) can be modified as follows: [51]

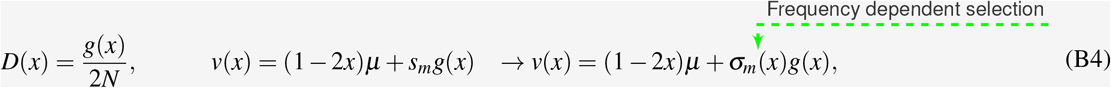

where *µ* is the mutation rate (assumed here to be symmetric for wild-type to mutant and vice versa), *s*_*m*_ is the mutant selection coefficient, and *g*(*x*) = *x*(1 − *x*). We have used the effective frequency-dependent selection coefficient to modify the first term in the FPK equation, replacing the selection coefficient *s*_*m*_ in *v*(*x*) with the effective frequency-dependent selection coefficient *σ*_*m*_(*x*). The constraints of moderately weak selection and mutation which allows the FPK description to accurately capture Wright-Fisher dynamics means we must have 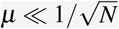 and 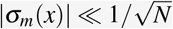 for all *x*, meaning 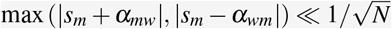.

Note that the FPK description reduces to a replicator-mutator equation in the limit as population size *N* → ∞, with the replicator portion exactly matching what we used above.

**The most likely evolutionary outcome as the mode of the probability density function**

The stationary solution *ρ*^eq^(*x*) for the FPK equation in **Eq**. (B3) is

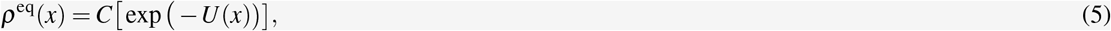

where *C* is a normalization constant, and 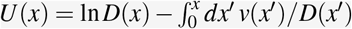.

The minima and maxima of the probability distribution can be determined by setting *∂ρ*^eq^ (*x*)/*∂x* = 0 (or equivalently *∂U* (*x*)/*∂x* = 0). In the no-interaction limit, *α*_*mw*_, *α*_*wm*_ → 0, for a population of size *N* with mutation rate *µ* > 1/2*N* and intrinsic selection coefficient *s*_*m*_, the mode of the stationary probability distribution (i.e. its global maximum) is given by the following relation:

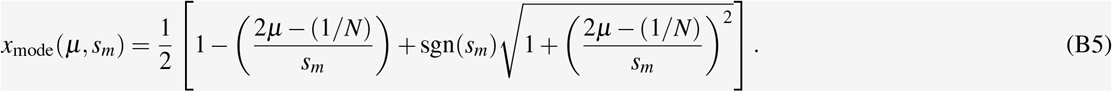

We use this modal population fraction, *x*_mode_, the peak of the stationary probability density function, to define equivalent regimes. See **Methods** and **Supplementary Information** for further details.

Real-world evolutionary dynamics (with or without interactions) are *stochastic*, with noise introduced through stochastic processes such as genetic drift. These sources of inherent noise, as well as measurement uncertainty, can significantly alter the recorded dynamics and steady states of the systems. It is unknown, from both the mathematical and experimental perspectives, how noise impacts our ability to distinguish interacting and non-interacting systems. To address this question, we extend our analysis to an FPK – a mathematical model of frequency-dependent evolution that includes stochasticity via genetic drift in addition to mutation. We use this model’s partial differential equation to capture the time evolution of the probability density function of the mutant fraction.

### Frequency-dependent selection in the Fokker-Planck-Kolmogorov model

Our reparametrization of the payoff matrix (**Eq**. (B1)) highlights how the payoff terms are interdependent in replicator systems and describes how the deterministic outcomes (game quadrants) depend on the interaction and selection terms. Further, we have outlined how the balance between the interaction and selection coefficients can result in maintenance, mirroring, masking, and mimicry (**Eq**. (2)). We wish to understand the possibility that these interaction classes occur in stochastic systems with mutation. To do so, we analyze how the probability density distribution of the mutant fraction, the solution to a partial differential equation, shifts with different interaction strengths. To define the four interaction classes, we first express the FPK stationary solution in terms of the selection and interaction terms (*s, α*_*wm*_, *α*_*mw*_). In particular, our concepts of maintenance, masking, mirroring, and mimicry rely on how the modal value of this solution shifts relative to the mode of the probability density distribution for a system *without* interactions.

In the modification of the population by inter-population interactions, our specific cases of interest reflect the position of the modal value of the population with respect to the initial modal value of a system without these interactions (**Eq**. (B5)). However, while numerical calculation of the modal value for given interaction coefficients and intrinsic selection is straightforward (see Supplementary Information), a general expression for the peak of the distribution with interactions is technically possible, though not particularly illuminating, as the solution of a cubic equation.

Fortunately, we can still easily analytically examine the conditions in which the modal value of an interacting system emulates the modal value for a non-interacting system. Consider the non-interacting modal frequency *x*_mode_(*s*^′^) (given by **Eq**. (B5)) for an arbitrary selection coefficient *s*^′^, and the effective frequency-dependent selection coefficient 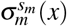 (given by **Eq**. (1)) for a given intrinsic selection coefficient *s*_*m*_ (and interaction parameters *α*_*mw*_ and *α*_*wm*_). The modal frequency with interactions will be the same as the modal frequency without interactions but with intrinsic selection coefficient *s*^′^ when

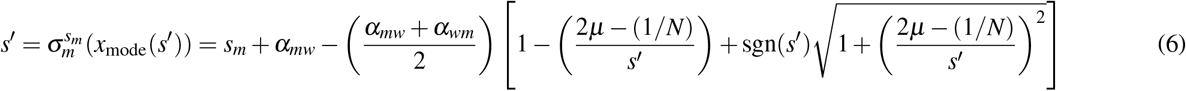

i.e. when the effective frequency-dependent selection coefficient *evaluated at the modal frequency for intrinsic selection s*^′^ is equal to *s*^′^, along with the additional condition

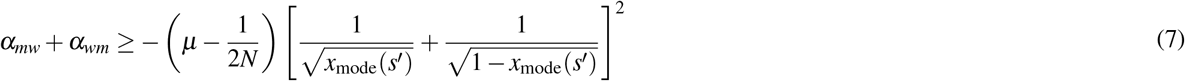

This second condition ensures that the probability for the interacting system has a unique maximum between 0 and 1, allowing it to properly emulate the non-interacting system (see Supplementary Information).

### Maintenance, masking, mimicry, and mirroring can occur in the presence of mutation and noise in the Fokker-Planck-Kolmogorov model of frequency-dependent evolution

To investigate the previously described four interaction classes, we can use our formula **Eq**. (6) constraining interaction parameters for emulation of non-interacting mutational, stochastic systems, with conditions summarised below in **Eq**. (8). Recall that the condition described in **Eq**. (7) above must also be applied.

SELECTION AT EQUILIBRIUM

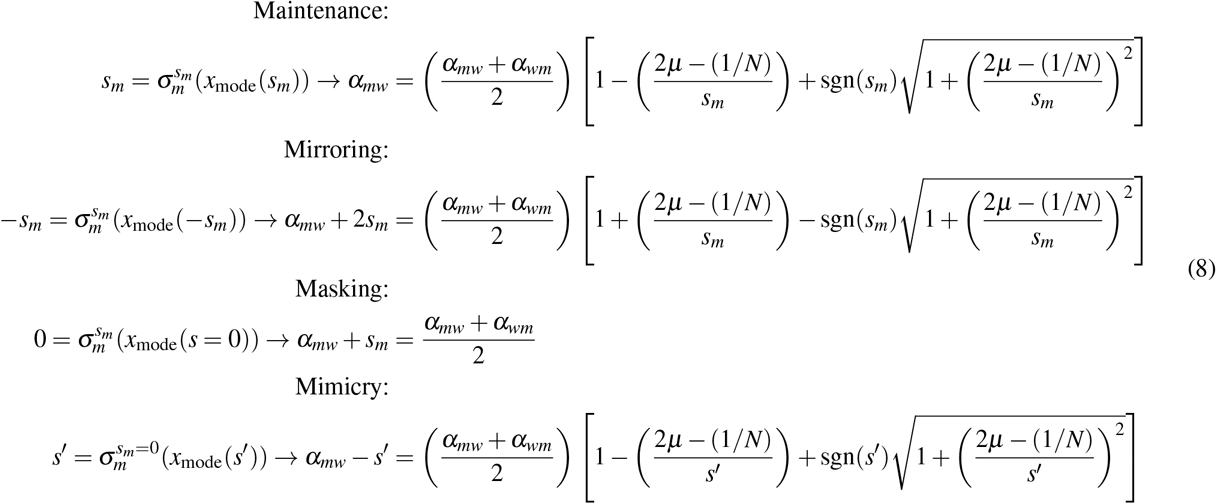

The surfaces described in **Eq**. (8) define the maintenance, masking, mimicry, and mirroring regimes. These regimes are defined by constraints on one interaction coefficient as a function of the other system parameters. Thus, each condition is a surface in three (for masking) or four-dimensional space (for the other regimes). The graphical representation of these relationships is illustrated for a range of fixed selection (*s*_*m*_) and mutation (*µ*) parameters in the Supplementary Information.

The Fokker-Planck-Kolmogorov (FPK) equation describes Wright-Fisher dynamics (in the large population limit, for relatively weak selection and mutation), and so to verify that our theoretical restrictions (on *α*_*wm*_ and *α*_*mw*_) in the FPK model result in the behavior discussed above, we simulated the dynamics of an evolving population with the original selection coefficient and then the specified interaction terms for the class behavior (**Fig 4**). The plots at the top of this figure display the theoretical relationships (**Eq**. (8)) for each example case. Initial simulations with no interactions (“Selection only”, meaning only *intrinsic* selection) are compared to simulations with added interaction terms for three different intrinsic selection coefficients and two mutation rates. Note that, while our analytical conditions relate to the modal value of distributions and the simulation results convey mean values of those distributions, these two values are negligibly different for distributions with a single maximum as the population size *N* becomes large, which is the situation here.

**Figure 4.**
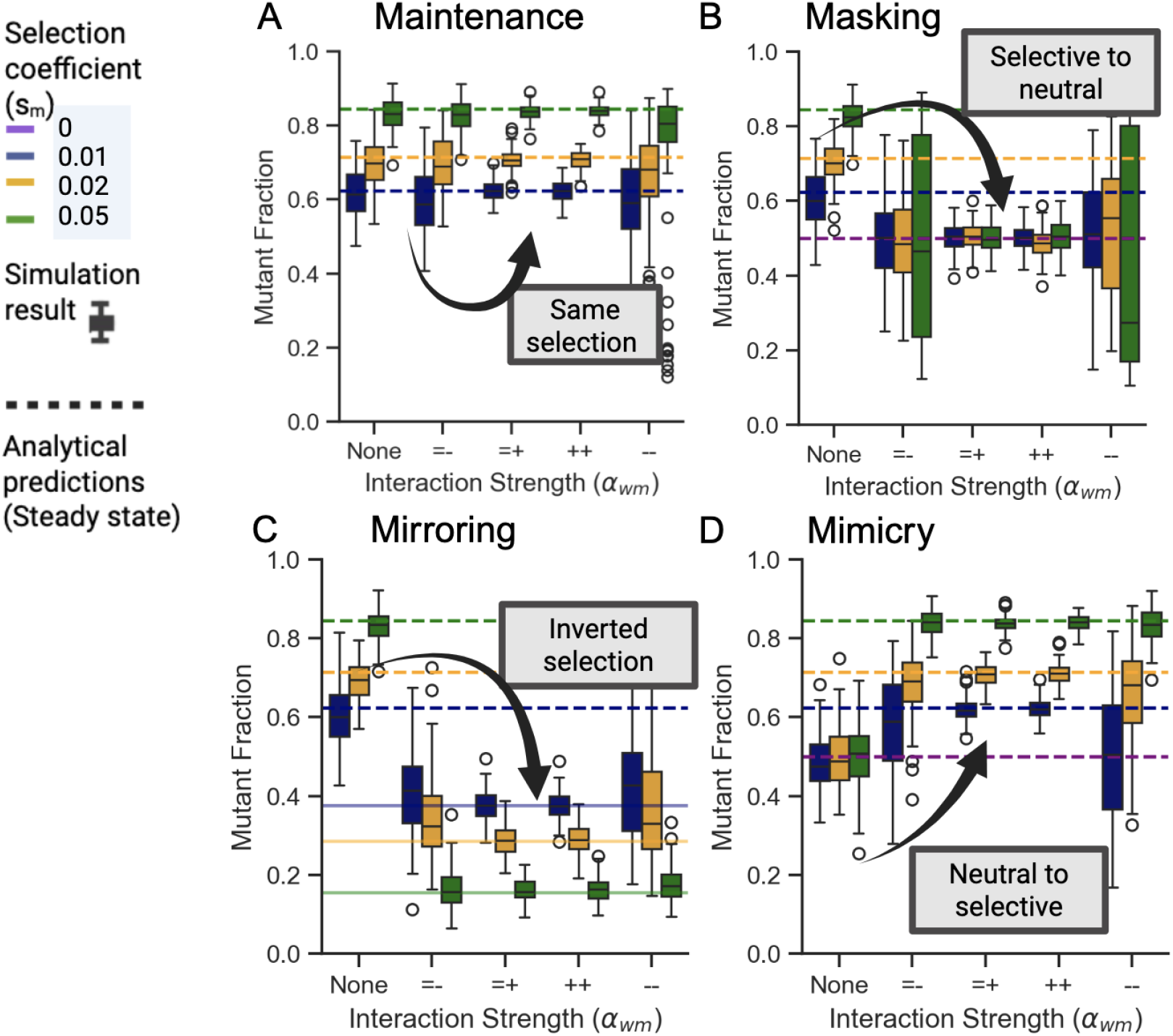
Analytical FPK conditions for the four interaction classes (Maintenance, Masking, Mirroring, and Mimicry) are validated in stochastic WF simulations. Wright-Fisher simulations were carried out for each scenario and a genetic population comprising individuals with one of two possible alleles (*N* = 1000), with no interactions (Selection only, i.e. *intrinsic* selection only), and with interactions. Dashed lines represent the theoretical modal value of the mutant fraction at different intrinsic selection values. For all scenarios, we compare simulations with no between-allele interactions (interaction coefficients →0), with simulations with increasing strength of *α*_*wm*_ interaction coefficient. Columns represent = −, lower limit of behavior meeting unique maxima condition, = +, upper limit of behavior meeting unique maxima condition, −−, less than unique maxima, ++ greater interaction strength than upper limit. The second coefficient, *α*_*mw*_, is derived using the relationships defined in **Eq**. (8). **(A)** For maintenance, the simulations maintain the same mean wild-type fraction as in the case with no interactions. **(B)** For the masking scenario, the interaction-free modal fraction of the wild type at each selection coefficient is not 50%. When interactions are introduced, the modal population changes to 50%, the equivalent of no selection. **(C)** In the case of mirroring, the population fraction for three different selection coefficients and interaction coefficients align with the analytical model. **(D)** In the mimicry case, the interaction-free population comprises 50% mutant as the mutant population has no intrinsic selection coefficient. With the mimicry interactions, we recapitulate the specific selection coefficients in each case. All simulations show agreement with our theoretical conditions across different strengths of *α*_*mw*_ and different values of *s*_*m*_ and *µ*. Modal population values are obtained for each simulation by averaging over generations 1001-2000. Each box plot represents the distribution of these modes over 100 independent simulations. Boxes are colored by selection coefficient. Initial conditions for all simulations were 500:500 wild-type (0) and mutant (1) individuals. Validity is most sensitive to initialized selection and interaction strengths meeting the conditions for a unique maxima **(Eq. B5)**.

In the *maintenance* case, the no-interaction simulation results fall on the theoretical modal fractions (dotted lines in **Fig 4**) for the appropriate selection (*s*_*m*_) as expected. As predicted theoretically, the populations in the *maintenance* interaction cases remain centered on the same theoretical value as in the no-interactions case (“Selection only” in **Fig 4**). Furthermore, varying the magnitude of frequency-dependent interactions maintains the intrinsic selection.

In the *masking* case, the non-interacting simulation populations align with the theoretical predictions for a given intrinsic fitness difference favoring the mutant. Under *masking*, this intrinsic selection difference is indeed nullified by frequency-dependent interactions. The mean simulated population value after *masking* is centered on the equal 50:50 split.

For *mirroring*, the no-interaction cases are aligned with the analytical mode expected for the given selective differences, yet the simulated populations under *mirroring* interactions are all perfectly reflected; the selection advantage of one population is inverted by the frequency-dependent interaction.

In the *mimicry* case, the theoretical modal values for all simulations with no interactions are at 50% mutant, as marked with the purple dotted line in **Fig 4**, and the simulated results without interactions match that value. However, the simulated populations with *mimicry* interactions center theoretical modes for three non-zero selection coefficients.

In summary, we derive a general expression for the equilibrium distribution of a population obeying the FPK equation with selection and arbitrary interactions. We reveal, using analytical theory and simulations, the potential impacts of game interactions and their ability to completely alter evolutionary dynamics. We find the critical boundaries at which these game dynamics either maintain a population dominated by the originally fitter genotype (maintenance), move a population from the domination of the fitter genotype in monoculture to the domination of the other (mirroring), promote domination of one subpopulation in the absence of intrinsic selection differences (mimicry), or even promote the heterogeneity of a population by leveling the playing field (masking).

### Interactions can produce maintenance, masking, and mirroring effects in *in vitro* experiments

To demonstrate how the reformulation of the payoff matrix can provide new frameworks to discuss how payoff interactions and dependencies produce certain experimental outcomes, we carried out a literature search to identify directly measured payoff matrices or co-culture experiments from which the EGT payoff matrix [21] can be defined (see Methods).

It is important to state that when using population growth rates to define payoffs in the replicator equation, we assume that the net growth rate is proportional to the number of individuals in the population (see Supplementary Information for derivation of the replicator equation).

The traditional evolutionary game theory representation of payoff matrix components plots the relative invasion fitnesses (*a*_21_ − *a*_11_, *a*_12_ − *a*_22_). We plot 20 published experiments in this space (**Fig 6A**). The color represents the publication from which the payoff matrix is obtained, and each dot is an individual reported measurement of the relative payoff (as determined directly from net sub-population growth rates). Numbered experiments represent different payoff results from the same paper and are numbered to highlight specific ordering and positional changes between plots. While this traditional plot is sufficient to determine the proportions of populations at equilibrium, the plot is non-unique with respect to the intrinsic and extrinsic contributions to the payoff matrix. We used our payoff matrix reformulation and examined published results with sufficient information to determine the magnitudes of selection and interaction effects. [21, 27, 28, 45, 46, 52] We show the interaction-selection plot, presenting the frequency-dependent interaction coefficients (*α*_*mw*_, *α*_*wm*_), on the axes, divided by the cell-intrinsic selection coefficient *s*_*m*_. This new interaction-selection plot illustrates the direction and intensity of the two interaction terms in a measured payoff matrix. Within this plot, the unit circle (blue) as shown in **Fig 6B** represents the boundary between dominance of frequency-dependent versus intrinsic selection effects. Within this circle, a system’s dynamics are dominated by intrinsic fitness differences. Outside of the circle, the dynamics are dominated by frequency-dependent effects. This visualization provides additional information, distinguishing strong and weak frequency-dependent effects relative to selection that result in very similar equilibrium proportions (i.e. points 1 and 6). This plot allows us to see the relative strengths of individual inter-population interaction-effects compared to selection. For example, payoff matrix **7**, with large positive *a*_21_ − *a*_11_ value, and large negative *a*_12_ − *a*_22_ value in **Fig 5A** lies close to the unit circle in **Fig 5B**. We see that for Farrokhian *et al*. [27](pink, 0-4), the relative strength of interactions is higher in 3 than 4 (**Fig 5B**) and the sign of the interaction effect on the mutant is reversed in the no-drug condition (0)(**Fig 5B**). We note that all of the bacterial interaction terms lie within [-2,2]. The most extreme examples of interaction-to-selection ratios 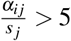 are in cancer models (e.g., 6: cell line [21]).

**Figure 5.**
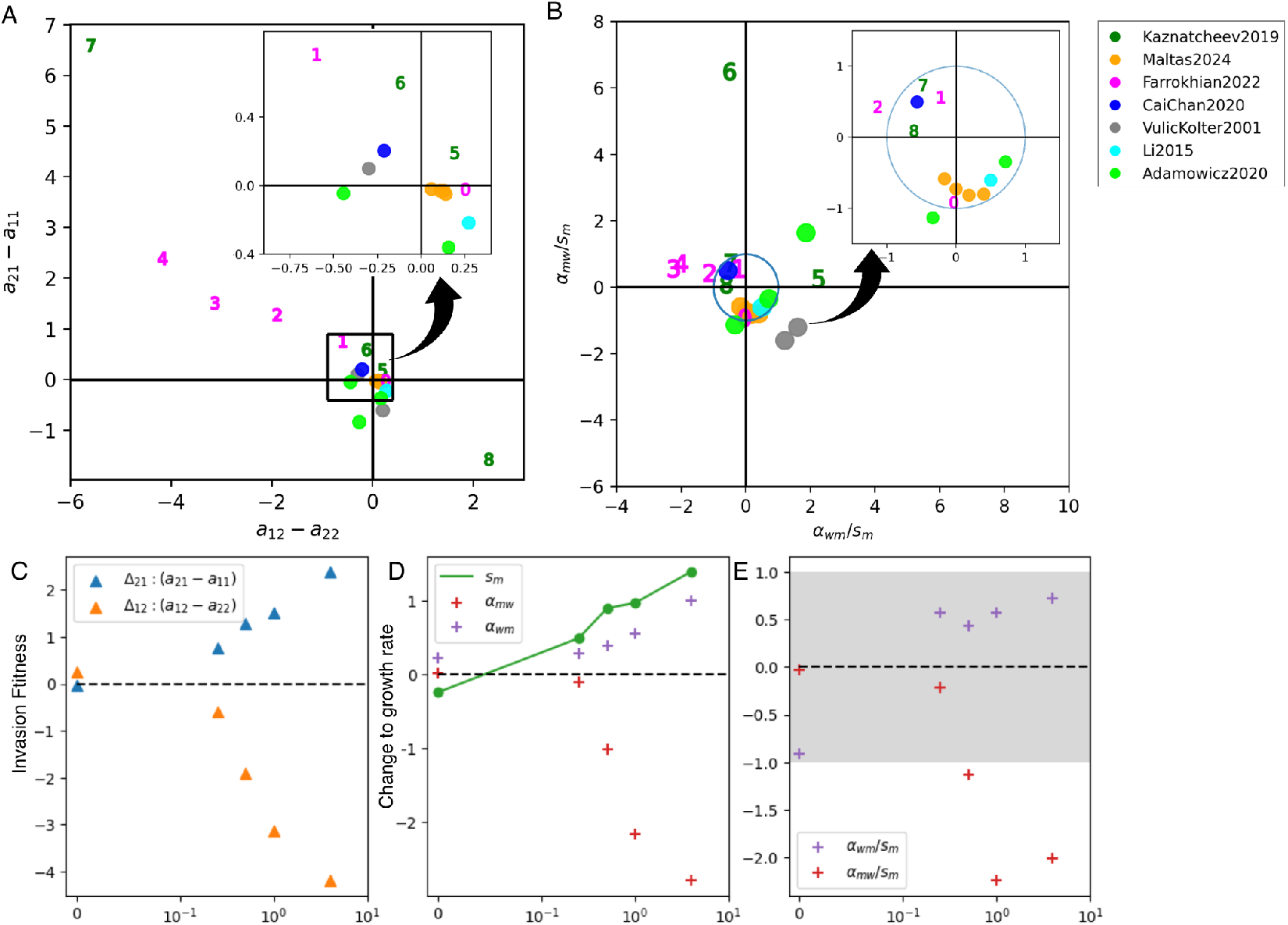
Interaction strengths decoupled from intrinsic selection show differential dominance of frequency-dependent interactions. Games, colored by publication reference, as measured in published experiments. Farrokhian (0-4) and Kaznatcheev (5-9) experiments are numbered for communication and illustration purposes. **(A)** Experimental measured games are displayed in “traditional” relative-growth-rate-encoded game space. The inset plot expands the area for visibility. **(B)** Interaction-selection plot. Distribution of published experimental interaction values relative to intrinsic selection. The unit circle (blue) is the boundary representing equal magnitudes of interactions and intrinsic selection. Experiments inside the circle are dominated by intrinsic selection, and interaction effects dominate those outside the circle. **(C)** The EGT invasion fitnesses from the payoff matrix (Δ_*mw*_ = *a*_*mw*_ − *a*_*ww*_, Δ_*wm*_ = *a*_*wm*_ − *a*_*mm*_) for increasing drug concentration (Farrokhian et al [27]). **(D)** The selection and interaction coefficients from the re-parameterization of the payoff matrix. The selection coefficient *s*_*m*_ is also plotted. **(E)** Interaction-selection ratio with changing drug concentration for Farrokhian et al. experiments (0-4) [11]. The shaded rectangle (vertical axis values < 1) corresponds to the region in which the selection coefficient is stronger than the interaction component.

**Figure 6.**
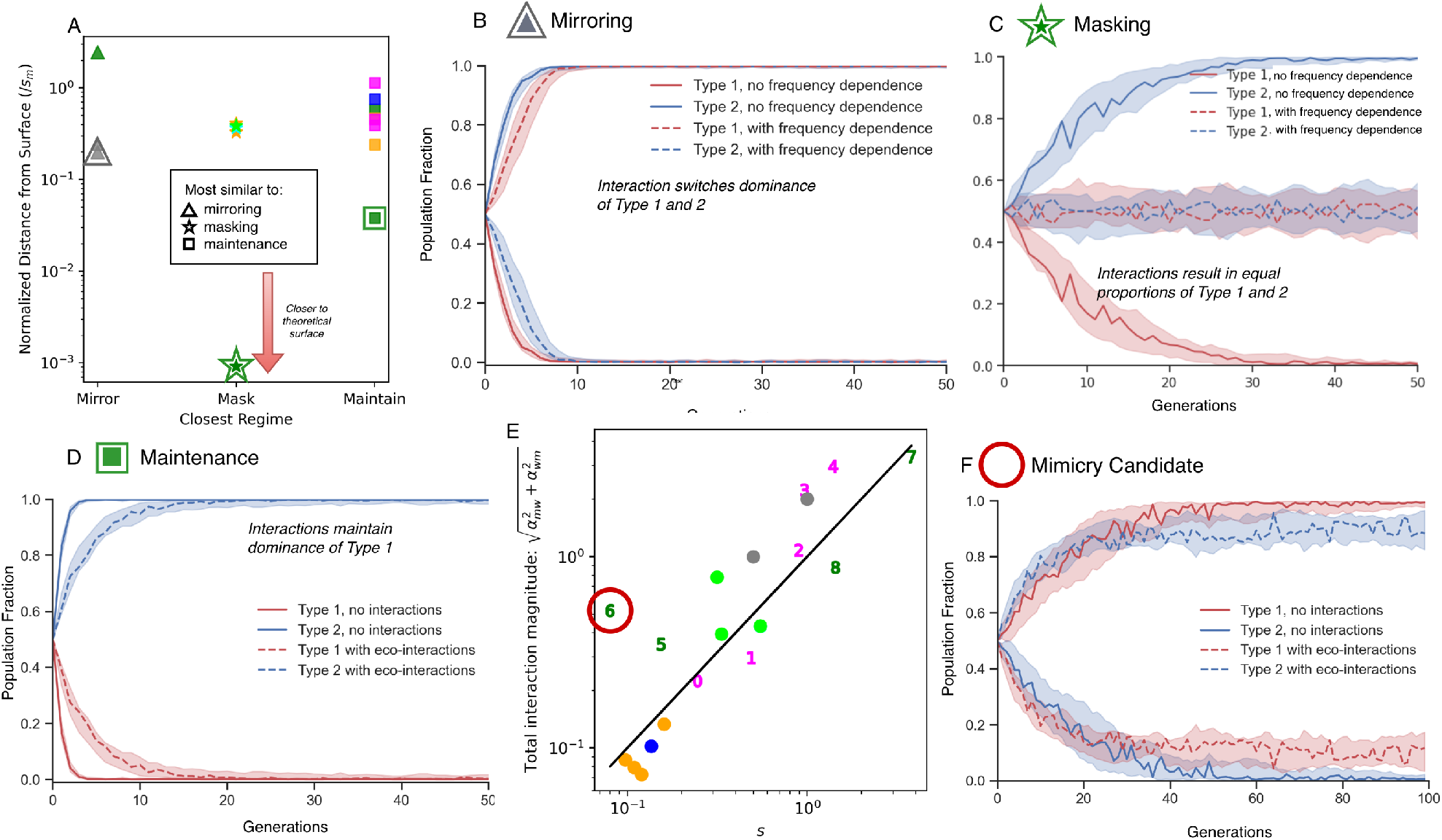
**(A)** Distances of published payoff composition from theoretical maintenance, masking, and mirroring surfaces are shown. Only experiments meeting the condition of having a single stationary point are included. The experimental payoff matrices closest to masking (green star), maintenance (blue square), and mirroring (pink triangle) are highlighted. **(B-D)** Simulations using the payoff matrices from the highlighted experiments in **(A)** illustrate the time-dependent frequency-dependent-evolutionary dynamics. The population dynamics are simulated for the three experiments: most mirror-like, masking-like, and maintenance-like. **(E)** Selection versus strength of interaction is plotted to identify the most likely candidate for mirroring. **(F)** Experimental payoff matrix values plotted with and without interaction components. While the intrinsic selection difference is small, the difference is large enough to create a selection difference.

An additional advantage of this new representation is its ability to distinguish qualitatively similar systems driven by different underlying mechanisms. In other words, two systems may occupy the same quadrant in the traditional game-theoretic space and therefore exhibit comparable equilibrium frequencies but may emerge from different underlying selection and interaction components. Our approach allows us to extract the changing relative magnitude of each component.

We explore this further by illustrating how the decomposition provides more intuitive information than the original payoff matrix and invasion fitnesses alone (**Fig 5C-E**). While prior work has incorporated drug-dependent modifications to payoff matrices, these effects are typically treated as undifferentiated net changes to the payoff entries. In contrast, our framework uniquely decomposes fitness into an intrinsic population-genetic selection term and an interaction-dependent component, allowing the intrinsic term itself to be modeled as drug-dependent and distinguished from interaction-mediated effects. [53] Here, the genotypic selection coefficient increases for the mutant with increasing drug concentration(**Fig 5(D)**). In the Farrohkian experiments, an increase in drug concentration results in an expected increase in the selective advantage of the mutant (*s*_*m*_). What is less intuitive is that a positive effect of the wildtype on the mutant increases proportionally (*α*_*wm*_) to this selection advantage, and the effect of the mutant on the wildtype becomes very negative (*α*_*mw*_).

We have previously presented the concepts of maintenance and masking to highlight that when frequency-dependent interaction effects between populations are present in co-culture, they can reinforce prior assumptions. Under the assumption that interactions or intrinsic selection processes are absent, one may overlook biologically relevant, potentially confounding factors that give rise to an equivalent aggregate outcome. To understand the extent to which specific regimes of maintenance, masking, and mirroring may be present in real multi-species systems, we examine the experimental coordinates of the selection and interaction coefficients and their position with respect to the aforementioned theoretical regimes.

We first applied the constraint (Eq. **??**) that for an experiment to be considered as emulating a non-interacting system, the payoff parameters must produce a single maximum in the stationary probability density distribution. To estimate proximity to maintenance and mirroring surfaces, we also approximated a fixed *µ* = 0.001 for co-cultured populations theoretically accessible to each other through mutation.

Each of the *maintenance, masking*, and *mirroring* regimes is defined by a function *α*_*wm*_ = *f* (*s*_*m*_, *α*_*mw*_, *µ*) (**Eq**. (8)). These conditions thus correspond to analytically defined manifolds. The minimum distance of the experimental system from each theoretical regime was calculated as the perpendicular distance from the measured value [*s*_*m*_, *a*_*mw*_, *a*_*wm*_] to the closest projection of the point onto the surface. These values are shown for each paper and each condition in **Fig 6A**. The distance from these theoretical surfaces is normalized by the selection coefficient *s*_*m*_. We note that three payoff matrices (**Supplementary**

**Table S2**: 2,3,12) had selection coefficients |*s*_*m*_| > 1. This breaks our weak selection assumptions used in Wright-Fisher and Fokker-Planck modeling.

Multiple systems are seen as approximating mirroring (triangle), masking (star), and maintenance (square), where experiments closer to the theoretical regimes are closer to the *x*-axis. The experiment most closely matched to each interaction class is highlighted. One of the experiments from Volic *et al*. (grey, triangle) is closest to mirroring, and two of the Kaznatcheev *et al*. experiments (green, star, square) are closest to masking and maintenance, respectively. In **Fig 6B-D** we simulate the frequency-dependent evolutionary dynamics of the populations using the published parameters for the highlighted experiments. Dynamics are plotted without the interaction components (solid lines) and then when the interactions are present (dashed lines). The Kaznatcheev experiment in **D** maintains approximately the same selection coefficient. There is increased noise in frequency-dependent dynamics, and some deviation from exact maintenance is noticeable on this time scale (0-50 generations). This is clear in contrast to the alignment to mirroring and masking dynamics in **B** and **C**.

*Mimicry* is more complex to examine in experiment. Although the presentation of a selective advantage from purely frequency-dependent means is an interesting evolutionary mode/adaptation, testing for it requires prior knowledge of the otherwise arbitrary “target” selection coefficient *σ*. As mimicry requires the intrinsic selection coefficient to be equal to zero, we highlight the experiment with the smallest selection coefficient relative to interaction strength (**Fig 6E**). In the simulation of these game properties (**Fig 6F**), the resultant dynamics do not produce clear mimicry behavior. Critically, there is not a neutrality of the non-interacting dynamics as defined for mimicry.

Key details about the publications and data points are shown in **Table 2**, and the complete data table is included in the **Supplementary Information** (**Table S2**). For each publication, either the directly reported payoff matrix or a payoff derived from the published growth rates in monoculture and coculture were used (see **Methods**). It is worth noting that the Cai et al. paper [45] uses metabolic models to determine a numerical payoff matrix.

**Table 1.** Key definitions.

| Term | Definition |
| --- | --- |
| Evolution | Temporal changes in the frequency of alleles within a larger population. |
| Fitness | The effective reproductive rate, growth rate (per capita) of an individual from a subpopulation. |
| Payoff | The components of the payoff matrix reflect the fitness of players of a given strategy in the presence of players of another given strategy. |
| Interaction | Effective change in reproductive fitness due to the presence of another strategy. The fitness as a result of self-interaction is indistinguishable from the isolated intrinsic growth rate in the non-density-dependent model. |

**Table 2.** Experimental payoff systems and associated papers. Results from the following papers were used to derive corresponding selection and interaction coefficients in their payoff matrices. *Li *et al*. found that the relationship between frequency and fitness was non-linear, but the linear assumption in our model was used for linear payoff matrix estimation from their data.

| Author | Year | System | Variable | Number of matrices |
| --- | --- | --- | --- | --- |
| Vulic and Kolter [52] | 2001 | Bacteria (*E. coli*) | Mutation | 2 |
| Li *et al.* * [19] | 2015 | Bacteria (*Curvibacter*, *Duganella*) | Species | 1 |
| Kaznatcheev *et al.* [21] | 2019 | Non-small cell lung cancer | Drug+CAFs | 4 |
| Adamowicz *et al.* [46] | 2020 | Bacteria (*E. coli* and *S. enterica*) | Mutation | 3 |
| Cai *et al.* [45] | 2020 | In-silico | Metabolic simulation | 1 |
| Farrokhian *et al.* [27] | 2020 | Non-small cell lung cancer | Mutation, Drug | 5 |
| Maltas *et al.* [28] | 2024 | Non-small cell lung cancer | Mutation, Drug | 4 |

## Discussion

The study of co-evolving populations often conceptualizes separate intrinsic and interaction contributions to fitness. However, the replicator equation does not, despite being commonly used to interpret and study biological evolution. [24] In this manuscript, we framed the payoff matrix from the replicator/replicator-mutator frameworks as consisting of intrinsic growth-term contributions with interactions as modifiers to this rate. This creates a novel interdependency, not present within other decompositions of the EGT payoff matrix, that we study further. [53] We used this framework to discuss the conditions under which the dynamics, or steady state, of an interacting population may be indistinguishable from the possible outcomes of non-interacting ones. We use this novel framing to directly compare the interaction-selection strengths in a new plot that can visualize and explore the relative strength of intrinsic selection versus interaction. In deriving analytical conditions for resultant system behaviors (maintenance, masking, mimicry, mirroring), we also show that a selection-normalized distance from each surface can be obtained, and we find that the ability for interactions in co-culture to emulate interaction-free dynamics depends on the mutation rate. The exploration of this distance and the stability and prevalence of these behaviors in the higher-order evolutionary landscape has yet to be explored.

Our work builds on existing experimental and mathematical descriptions of co-evolutionary dynamics. [27, 40–42, 54, 55] By reframing the EGT payoff matrix such that interactions and monoculture fitnesses are separated, we come to a form that can be easily modeled using a frequency-dependent form of the diffusion-drift (FPK) equation, as derived in population genetics. [43, 44, 51] Separately, this mapping also allows us to observe that dilemma strength scaling [54] becomes a scaling proportional to the selection coefficient. Although not explored here, incorporating the selection coefficient directly into the payoff matrix also enables the future incorporation of drug-dependent selection models (e.g., Hill-type dynamics) into the payoff. This would allow fitting the selection term across game measurements to dose-response curves and modeling how systems can traverse “game space” with and without proportional changes to off-diagonal interaction terms.

We, as others have before us, reinforce that growth dynamics in mixed populations may readily bear little resemblance to monoculture growth rates, and that initial seeding (e.g., experimental or metastatic) ratios may have strong impacts on resultant steady states. Our framework also highlights the critical but potentially invisible nature of between-population interactions in an experiment. We show how the presence and strength of game interactions, even of the same order of magnitude as the underlying selection coefficient, have the potential to modify the steady state of an evolving population to many outcomes. This range includes outcomes producible by a frequency-independent system. Our ideas find real-world importance in the ongoing study of human diseases, where exciting *in vitro* work and retrospective modeling of clinical data continue to provide empirical evidence for the presence of frequency dependence in mixed populations. [56–58] Experimental scientists using “game assays” have also begun to quantify these interactions between cells in culture. [21] We use game theoretical concepts to model and explore the possible impacts of these interactions as they become increasingly important in the disease and cell-biology perspectives. [58, 59] Despite this growing experimental evidence for frequency-dependent interactions in microbial and cancer systems, most evolutionary studies still treat fitness as a fixed, frequency-independent property. Without specific techniques designed to robustly assay frequency-dependent game interactions, [21, 27], the magnitude and impact of interactions cannot be quantified. Neglecting interaction effects has widespread potential ramifications, for example, when assaying chemotherapeutic drugs in isolated cell lines, when developing cancer cell lines *ex vivo*, and when interpreting evidence for neutral evolution in tumors. [60–64] We reviewed the experimental literature for payoff matrices, bringing together a range of data in a single framework via our novel interaction-selection plot that compares effects. Further work beyond the scope of this study is required to understand whether these modes of behavior can be defined in systems exhibiting non-logistic growth or coupling with the environment. [23, 65] A promising area for cross-evaluation includes integrating findings from density-dependent modeling of overlapping datasets. [66]

Having shown that deterministic game theory is significantly limited in distinguishing the outcomes of interacting and non-interacting systems, we have only begun to explore the potential to distinguish these outcomes within stochastic frameworks. While we show that our stochastic analytical and simulation results are in agreement, we have not approached the reverse problem, the inference of game parameters from the simulation results. This is an important next step, as this stochastic description of co-evolutionary dynamics over time may allow us to leverage larger quantities of inherently stochastic experimental results to infer interaction properties. The goodness of fit of this model to experimental or clinical data could also be compared to alternative frameworks, as has been laid out with other models in recent work. [58, 66]

Further limitations of our modeling approach include the non-mechanistic formulation of game interactions and the assumed linearity of the frequency-dependent interactions, an assumption that likely does not hold across all systems. Additionally, to analyze and discuss previously published results from the literature, simplifying assumptions were made about mutation rates. Although in the published results we see several relatively strong frequency-dependent effects, the results shown are not exhaustive. And although interactions here are evident, there is likely a positive bias in the literature toward the publication of experimental results where interactions are present. Whilst our work focused on the modal value of the equilibrium distribution after a long period, the varying higher-order properties of the FPK distributions in systems with games may also have altered evolutionary dynamics over time. This has been noted in previous work, [37], where Kaznatcheev demonstrated that evolution is, in some cases, accelerated by game interactions. Future work could derive explicit expressions for signatures of game dynamics that are encapsulated in the higher-order properties (e.g., standard deviation, skew) of the equilibrium solutions. Identifying these factors may be critical in interpreting the existence and strength of cell-cell interactions in experimental populations. Our work is limited to the study of two populations/alleles, and broadening this framework and concept to populations with more than two genotypes may not be analytically solvable, but could be approached with numerical techniques.

Decomposition of the payoff matrix provides a biologically meaningful formulation of the payoff matrix and the ability to independently modify the cell-intrinsic and frequency-dependent interaction contributions to the growth rate. In addition to supplying a new modeling paradigm and analytical solution in the case of added noise, this formalism provides an ideal starting point for deeper analysis of experimental results in the context of intrinsic selection and stochasticity. This model framework is also directly compatible with the integrated modeling of evolutionary therapy, mapping between monoculture and co-culture results in a stochastic mathematical description of an evolving population. [6, 47] These frameworks can assist calculations for optimal stochastic control and adaptive therapy in the presence of experimentally motivated drug and micro-environmental-dependent cell interactions. [27, 48, 58, 67–70]

## Materials and Methods

### Theory

To develop our initial theoretical approach, we build on a simple model of evolutionary game dynamics in the absence of noise and mutation. Game theory, in general, is the study of the dynamics that result from the interaction of different strategies played against each other. Specific strategies result in expected payoffs for the players, and thus the average payoff depends upon the frequency of strategies in the system. Differential game theory can describe deterministic solutions for these dynamics using differential equations. We assume symmetry in which the payoffs in a 2-strategy symmetric differential game can be presented as a payoff matrix in the following way [25, 71]:

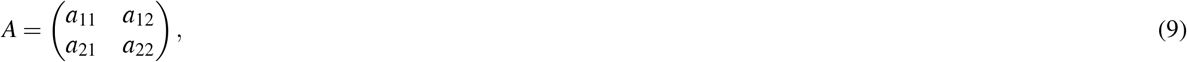

where *a*_11_ is the payoff for a player with strategy 1 playing against another player of type 1, *a*_12_ is the payoff for a player with strategy 1 playing against a player of type 2, *a*_22_ is the payoff for a player with strategy 2 playing against a player of type 1, and *a*_22_ is the payoff for a player with strategy 2 playing against a player of type 2. These values determine the expected payoffs (or “fitnesses” in the evolutionary case) for each player when the frequencies of strategies in the population are known.

In evolutionary models, available “strategies” equate to the cell type as a cell does not choose, but inherits its strategy; the strategy (phenotype) and identity (genotype) are assumed to form a one-to-one mapping. Under this assumption that a cell’s genome determines its strategy, we use a simplified model comprising a single locus with two alleles, the wild-type and the mutant, the latter harboring a resistance mutation at the site of interest. In genetic population models, strategy proportions change when cells undergo self-replication, with fitter strategies reproducing at a faster rate.

The underlying assumption of replicator dynamics – that the growth rate of type *i* is proportional to the number of cells of type *i*, with proportionality called the fitness in the evolutionary case – leads to frequency dynamics depending on the difference between the fitness of type *i* and the average fitness over all types. This formalism and accompanying equations are referred to as replicator dynamics. [24, 71] In the 2-dimensional case with payoff matrix *A* from Eq. (9), the replicator equation takes the following form:

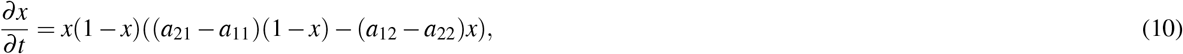

where the proportion of type 2 (mutant) is *x*, and thus the proportion of type 1 (wild type) is 1 − *x*. Without the addition of noise, the replicator equation gives us deterministic solutions for the evolutionarily stable strategies present. [72] Without mutation in the population, this equation has stationary solutions at *x* = 0, *x* = 1, and *x* = (*a*_21_ − *a*_11_)/((*a*_12_ − *a*_22_) +(*a*_21_ − *a*_11_)). Following this, different relative magnitudes of *a*_11_, *a*_12_, *a*_21_, and *a*_22_ result in various evolutionarily stable solutions. One can construct the 2-dimensional “game space”, which has the axes *a*_21_ − *a*_11_ and *a*_12_ − *a*_22_ as shown in **Fig 2**. In the top left *a*_21_ > *a*_11_ and *a*_22_ > *a*_12_, the wild-type always outgrows the mutant. In the bottom right, the converse is true. In the top right, *a*_21_ > *a*_11_ but *a*_12_ > *a*_22_, so the species co-exist. In the bottom left, *a*_21_ < *a*_11_ and *a*_12_ < *a*_22_, there is an unstable mixed equilibrium that cannot persist within a stochastic system. A specific two-player game payoff matrix has a position in one of these quadrants that categorizes it into a certain universality class. Games in the same quadrant generate similar dynamics. Different specific games and different quadrants are named across the EGT and ecology literature, e.g., Prisoner’s Dilemma (top left), Snow Drift (bottom left), Stag Hunt (bottom right), and Harmony (top right). The relationship between the classes of games in ecology versus EGT has been discussed previously. [55]

The derivations for the deterministic conditions of the different regimes are outlined in the **Supplementary Information**, as well as the derivation for the conditions in stochastic systems.

### Simulations

Using Python (version 3.11.5), we developed a stochastic Wright-Fisher type population model with discrete non-overlapping generations modified from Bedford *et al*. [73]. Creating a two-genotype model, we incorporated generational sampling that was weighted by frequency in ratios given by the coefficients of a payoff matrix. We used this to observe the simulated evolutionary trajectories and the stationary distribution before and after the addition of game interactions of different strengths. In particular, we asked whether our simulation results are consistent with game interactions predicted theoretically to maintain, mimic, mirror, or mask the expected evolutionary outcome that would be based on monoculture fitness alone.

The simulations (parameter ranges in **Table 3**), involved a constant population of size *N* comprised of two species, denoted wild-type (‘0’) and mutant (‘1’), undergoing mutation (rate *µ*) and selection at each generation. The sampling frequency of each population was based on the frequency-dependent fitness of each population calculated at each generation. The constant population size *N*, mutation rate *µ*, and mutant advantage *s*_*m*_ were all fixed for each simulation. When random net payoff matrix values or interaction strengths (*α*_*ij*_) were required, they were generated using a random uniform distribution within a given range. In each simulation, populations were allowed to evolve from an initial fraction *x*_*i*_ for 1000 generations, at which point the population fraction from each simulation was extracted.

**Table 3.** Methodological parameter values and ranges for the frequency-dependent Wright-Fisher model.

| Parameter | Symbol | Value(s) |
| --- | --- | --- |
| Mutation rate (per generation) | $\mu$ | $5 \times 10^{-2}, 1 \times 10^{-3}$ |
| Population size | $N$ | 1000-10000 |
| Generations | $t$ | 500-2000 |
| Initial fraction (i) | $x_i$ | (0,1) |
| Intrinsic selection | $s$ | (0,0.05) |
| Interaction effect of $i$ on $j$ | $\alpha_{ij}$ | (-0.2,0.2) |

### Intrinsic selection and extrinsic interaction values from the literature

We found co-culture experiments published in the literature, [21, 27, 28, 46, 52] and converted them to normalized and decoupled payoff matrices. Published payoff matrices that were derived by fitting a Lotka-Volterra model were excluded to avoid double-counting the intrinsic growth rate. [74] Decoupled payoff matrices are expressed in terms of both intrinsic selection and interaction growth rate effects (see **Eq**. (B1)). Our search was restricted to published results that explicitly reported experimentally derived payoff matrices [21, 27, 28]or publications with experimental growth rates for populations in both mono- and co-culture. Details of the publications, system or experiment type, and number of payoff matrices are shown in **Table S1**.

To calculate the payoff matrix and thus interaction values and selection from mono- and co-culture growth rate information, it is possible to construct an interaction (*growth-frequency*) plot for both populations (see Supplementary Information), from which the intersections (payoff values) can be inferred. With the payoff matrix generated, intrinsic and extrinsic growth rate modifiers can be calculated.

### Distance of experiments from maintenance, masking, and mirroring surfaces

The maintenance, masking, and mirroring surfaces were defined in **Eq**. (8). To calculate the proximity of the published experiments’ payoff values to the theoretical manifolds representing maintenance, masking, and mirroring (see GitHub for code), we used numerical optimization. Where mutation was plausible, a small mutation rate was assumed (*µ* = 0.001) in order to derive approximate distance from effect surfaces.

The unitless Euclidean distance between two points was calculated and minimized with respect to points on the surface to approximate the distance of the measured point from the respective manifold. Numerical optimization (numPy) was used to find the shortest distance between the measured value 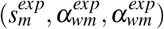 and points on the surfaces defined for maintenance, masking, and mirroring. A Euclidean distance function was minimized subject to this surface constraint. The optimized point is the projection of the original point onto the surface. Then the distance (*d*_*min*_) between the experimental point and its projection is calculated numerically. An illustration of this idea in less than four dimensions is in the Supplementary Information.

## Acknowledgments

We acknowledge Dr. Arda Durmaz, Dr. Arvid Agren, and Dr. Kyle Card for their helpful insight during the completion of this work.

## Author contributions

The following contributions were made by the authors.

Conceptualization - RBC, JG, MH, JS, DT, MS, JM.

Data curation - RBC.

Formal analysis - RBC, JG, JAP, MH.

Funding acquisition - JS.

Investigation - RBC, JG, JAP, MH.

Methodology - RBC, JG, MH.

Software - RBC, JG, MH, SL.

Supervision - JS, MH.

Validation - SL.

Visualization - RBC.

Writing – original draft - RBC, JG, MH.

Writing – review & editing - RBC, JG, JAP, MH, MS, SL, JS.

## Code and data availability

All of the Mathematica and Python scripts in this project can be found on GitHub at; https://github.com/rbarkerclarke/MaskMimicMaintain. The processed previously published experimental data is available in the supplementary files and the same repository.

