## Supplementary Information for "Hidden Interactions: Frequency-Dependence Emulates Selection-Driven Dynamics in Evolving Populations"

### S1 Transformation of the payoff matrix as it relates to the game space

The dynamics in a two-player game are often displayed as quadrants in a 2D “game space" with quadrants corresponding to different dynamical outcomes (**Fig. 5A**) [1]. Our variable transformation of the payoff matrix allows us to highlight how the quadrants are regions defined by interaction strengths relative to intrinsic selection and their analytical forms. In the intrinsic and extrinsic form of the payoff matrix, these axes correspond to $(g_m + \beta_{mw} - g_w)$ and $(g_w + \beta_{wm} - g_m)$. Here $g_i$ is the intrinsic growth rate of type $i$ and $\beta_{ij}$ is the maximum extent to which interaction with type $j$ affects the growth of $i$. In this transformed basis we can also write these as $g_w(\alpha_{mw} - s_m)$ and $g_w(s_m + \alpha_{wm})$ where the quadrants can be defined by conditions on $\alpha_{mw}$ and $\alpha_{wm}$ relative to the homogeneous mutant population selection coefficient $s_m$ (**Fig. 5B**). For the novel “interaction-selection" plot (**Fig. 5C**), we can plot the selection normalized interaction coefficients $(\alpha_{ij}/s_i,\ \alpha_{ji}/s_i)$. As we move through different values of $\alpha_{wm}$ and $\alpha_{mw}$ relative to $s_m$ the resultant equilibrium state is modified. We highlight the unit circle in this plot as a key region where intrinsic selection differences are dominant. As might be expected, within this circle, the game dynamics of the system, dominated by intrinsic selection, will align with the quadrant representing dominance by the same population. Deterministic boundaries between universality classes in all phase spaces occur along the lines $\alpha_{wm} = -s_m$ and $\alpha_{mw} = s_m$ For example, the cooperation quadrant (top right) is defined by $\alpha_{wm} > s_m$ and $\alpha_{mw} > -s_m$.

### S2 Deterministic Replicator Equation

The replicator equation (RE) describes the deterministic evolution of the frequencies of sub-populations, where the rate of change is proportional to the difference between the fitness of the subpopulation and the average population fitness.

**Derivation** Consider a population where the number of individuals using strategy $i$ is $N_i$, with total population $N = \sum_{j=1}^{n} N_j$. The frequency is

$$x_i = \frac{N_i}{N},$$

and the growth of each subpopulation is **assumed to be** proportional to its fitness:

$$\dot{N}_i = f_i N_i.$$

Subsequently, the derivative of the frequency is

$$\dot{x}_i = \frac{d}{dt}\left(\frac{N_i}{N}\right) = \frac{\dot{N}_i N - N_i \dot{N}}{N^2}.$$

If we substitute $\dot{N}_i = f_i N_i$ and $\dot{N} = \sum_j \dot{N}_j = \sum_j f_j N_j = N\bar{f}$:

$$\dot{x}_i = \frac{f_i N_i N - N_i(N\bar{f})}{N^2} = \frac{N_i}{N}(f_i - \bar{f}) = x_i(f_i - \bar{f}).$$

Therefore,

$$\dot{x}_i = x_i(f_i(\mathbf{x}) - \bar{f}),$$

where $\bar{f}$ is the average fitness of the population,

$$\bar{f} = \sum_{j=1}^{n} x_j f_j(\mathbf{x}). \tag{S1}$$

### S3 Derivation of the deterministic frequency-dependent selection coefficient

To derive the frequency-dependent selection coefficient for our deterministic model, we use the payoff matrix and fitness vector to calculate the selection coefficient. Our decomposition of the payoff matrix into selection and interaction components results in the following fitness vector:

$$
\begin{aligned}
f(\mathbf{x}) &= A\mathbf{x} \\
&= \begin{pmatrix} 1 & 1+\alpha_{wm} \\ 1+s_m+\alpha_{mw} & 1+s_m \end{pmatrix} \begin{pmatrix} x \\ 1-x \end{pmatrix} \\
&= \begin{pmatrix} 1+\alpha_{wm}x \\ 1+s_m+\alpha_{mw}(1-x) \end{pmatrix}
\end{aligned} \tag{S2}
$$

where $\mathbf{x} = (x_w,\ x_m)$ is the vector of frequencies of the wild-type and the mutant, where $\sum x_i = 1$ and can therefore be rewritten, $x = x_m = 1 - x_w$. Any component of the fitness vector, $f_i$, is given by $(A\mathbf{x})_i$. The first component of $f(\mathbf{x}) = (f_w(\mathbf{x}), f_m(\mathbf{x}))$ is the wild-type fitness $f_w(\mathbf{x})$, and the second component is the mutant fitness $f_m(\mathbf{x})$. The frequency-dependent effective selection coefficient $\sigma_m(\mathbf{x})$ that is to replace the constant selection coefficient $s_m$ turns out to be the difference between frequency-dependent fitnesses, divided by the frequency-independent wild-type fitness (which is set to 1):

$$
\sigma_m(\mathbf{x}) := \frac{f_m(\mathbf{x}) - f_w(\mathbf{x})}{f_w} = f_m(\mathbf{x}) - f_w(\mathbf{x})
$$

substituting the fitnesses in the 1D case as defined by the above payoff matrix, we obtain:

$$
\sigma_m(\mathbf{x}) = 1 + s_m + \alpha_{mw}(1-x) - (1+\alpha_{wm}x) = s_m + \alpha_{mw} - (\alpha_{wm} + \alpha_{mw})x.
$$

### S4 Replicator-mutator equation

The replicator-mutator equation (RME) [2,3] extends the RE to add mutation between the subpopulations. In a system with mutation, at each generation, mutations from type $i \to j$ occur at a rate $\mu_{ij}$ per unit time. There are multiple formulations of the RME, with small differences partially depending upon the order in which selection and mutation occur. We consider the following form, which matches the noiseless limit of Wright-Fisher dynamics for relatively weak selection and mutation (in this case, the order of selection and mutation does not matter):

$$
\dot{x}_i = x_i(f_i(\mathbf{x}) - \bar{f}) + \sum_{j \neq i} (\mu_{ji}x_j - \mu_{ij}x_i) \tag{S3}
$$

where the additional terms reflect the flow in and out of genotype $i$ due to mutation. In the limit $N \to \infty$, the Fokker-Planck-Kolmogorov equation (which is the high population and weak selection/mutation limit of Wright-Fisher evolution) reduces to this form of the replicator-mutator equation.

The transformation of the payoff matrix is used to calculate $f_i$ and $\bar{f}$, and we assume a symmetric mutation rate $\mu$. In this framework, the equilibrium solutions for the allele fraction, $x$, are the solutions to the following cubic equation.

$$
\dot{x} = \mu + (\alpha_{wm} - 2\mu + s_m)x + (-\alpha_{mw} - 2\alpha_{wm} - s_m)x^2 + (\alpha_{mw} + \alpha_{wm})x^3 = 0 \tag{S4}
$$

When interactions are nonzero, the solutions are functions of $\alpha_{ij}, \mu$, and $s_m$. For the interaction-free case where $\alpha_{mw} = \alpha_{wm} = 0$, this cubic reduces to a quadratic equation,

$$
\dot{x} = \mu + (s_m - 2\mu)x - s_m x^2 = 0 \Rightarrow x(t \to \infty) = \frac{1}{2}\left[1 - \left(\frac{2\mu}{s_m}\right) + \operatorname{sgn}(s_m)\sqrt{1 + \left(\frac{2\mu}{s_m}\right)^2}\,\right] \tag{S5}
$$

We can solve the cubic equation numerically for specific values of the interaction and selection coefficients, and impose equivalence of the interaction-dependent and interaction-independent equilibrium states. These numerical solutions can be used to illustrate the concepts of maintenance, masking, mirroring, and mimicry. In the presence of mutation, non-trivial solutions for maintenance emerge. Examples of maintenance, masking, mirroring, and mimicry, using numerical solutions, are demonstrated in **Figure S1**.

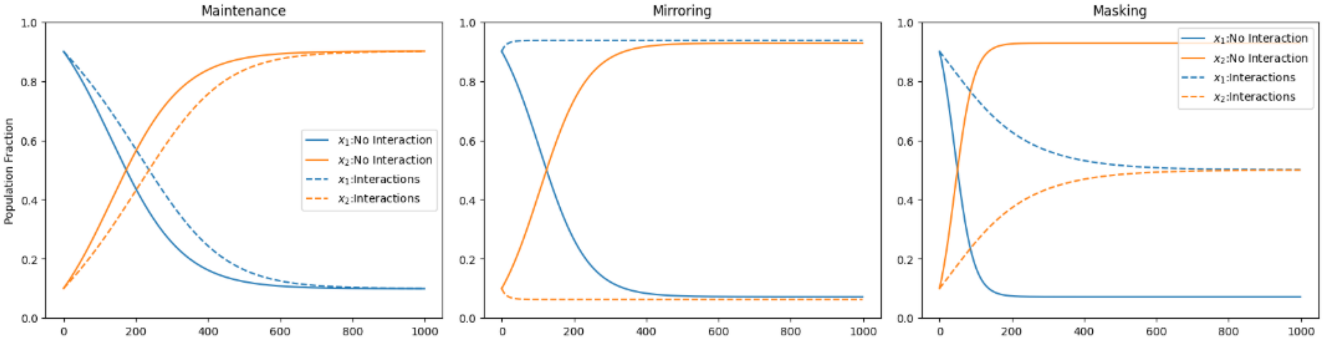


**Figure S1.** Maintenance, Mirroring, and Masking for the replicator mutator equation. Numerical solutions for deterministic dynamics shown with $\alpha_{mw} = \alpha_{wm} = 0$ (No Interaction) compared to $\alpha$ values determined by the conditions for equivalence between interacting and interaction-free systems. All examples have initial fraction $\mathbf{x}_0$ = [0.9,0.1], mutation rate $\mu = 0.01$ and $s_m = 0.15$.

### S5 Derivations of behaviors in the deterministic replicator equation

Maintenance, masking, mirroring, and mimicry behaviors of a frequency-dependent system are defined in comparison to a frequency-independent system with selection coefficient, $s_m$. Equivalence in behavior can be defined by the selection coefficient or the stationary solution(s). We use our reformulation to produce the following derivations for maintenance, mirroring, masking, and mimicry.

#### Regimes by identical dynamics

Our four regimes correspond to the equivalence between frequency-dependent system dynamics and frequency-independent ones. In the absence of mutation and drift, the selection coefficient determines the entirety of the evolutionary dynamics, and the behaviors can be defined by identical *dynamics* when the frequency-dependent selection coefficient is equal to the frequency-independent reference values:

##### Maintenance

$$\begin{aligned} \sigma_m(x) &= s_m \\ s_m + \alpha_{mw} - (\alpha_{wm} + \alpha_{mw})x &= s_m \\ \alpha_{mw} - (\alpha_{wm} + \alpha_{mw})x &= 0 \\ \Rightarrow \alpha_{mw} &= -\alpha_{wm} = 0 \end{aligned} \tag{S6}$$

##### Mirroring

$$\begin{aligned} \sigma_m(x) &= -s_m \\ s_m + \alpha_{mw} - (\alpha_{wm} + \alpha_{mw})x &= -s_m \\ \alpha_{mw} - (\alpha_{wm} + \alpha_{mw})x &= -2s_m \\ \Rightarrow \alpha_{mw} &= -\alpha_{wm} = -2s_m \end{aligned} \tag{S7}$$

##### Masking

$$\begin{aligned} \sigma_m(x) &= 0 \\ s_m + \alpha_{mw} - (\alpha_{wm} + \alpha_{mw})x &= 0 \\ \alpha_{mw} - (\alpha_{wm} + \alpha_{mw})x &= -s_m \\ \Rightarrow \alpha_{mw} &= -\alpha_{wm} = -s_m \end{aligned} \tag{S8}$$

##### Mimicry

$$\begin{aligned} \sigma_m(x, s_m = 0) &= s' \\ \alpha_{mw} - (\alpha_{wm} + \alpha_{mw})x &= s' \\ \Rightarrow \alpha_{mw} &= -\alpha_{wm} = s' \end{aligned} \tag{S9}$$

#### Regimes by steady state only

The deterministic replicator equation describing the non-neutral evolution of the subpopulation frequencies has either 2 or 3 equilibrium solutions, corresponding to deterministic steady states of the mutant allele fraction $x$. (Deterministic, mutation-free, neutral evolution – i.e. with no selection – is trivially stationary at its initial condition; no dynamics occurs.)

For the non-interacting $(\alpha_{wm}, \alpha_{mw} = 0)$, non-mutating, non-neutral system, there are two fixed points, at $x = 0$ or $x = 1$, where the stability depends on the sign of $s_m$ ($s_m > 0$ stabilizes $x = 1$, $s_m < 0$ stabilizes $x = 0$).

For the interacting case, there is additionally the third stationary point at

$$x^* = \frac{s_m + \alpha_{mw}}{\alpha_{mw} + \alpha_{wm}} \tag{S10}$$

In order for true emulation of non-interacting systems to occur, this stationary state cannot be a physically accessible state that differs from possible stationary states for non-interacting systems, thus we must have $x^* \leq 0$ or $x^* \geq 1$. These can be combined as:

$$\begin{aligned}
\left(x^* - \frac{1}{2}\right)^2 &\geq \left(\frac{1}{2}\right)^2 \\
\Rightarrow \left(\frac{s_m + \alpha_{mw}}{\alpha_{mw} + \alpha_{wm}} - \frac{1}{2}\right)^2 &\geq \left(\frac{1}{2}\right)^2 \\
\Rightarrow \left[s_m + \alpha_{mw} - \frac{\alpha_{mw} + \alpha_{wm}}{2}\right]^2 &\geq \left(\frac{\alpha_{mw} + \alpha_{wm}}{2}\right)^2 \\
\Rightarrow (s_m + \alpha_{mw})^2 - (s_m + \alpha_{mw})(\alpha_{mw} + \alpha_{wm}) + \left(\frac{\alpha_{mw} + \alpha_{wm}}{2}\right)^2 &\geq \left(\frac{\alpha_{mw} + \alpha_{wm}}{2}\right)^2 \\
\Rightarrow (s_m + \alpha_{mw})^2 - (s_m + \alpha_{mw})(\alpha_{mw} + \alpha_{wm}) &\geq 0 \\
\Rightarrow (s_m + \alpha_{mw})(s_m - \alpha_{wm}) &\geq 0
\end{aligned} \tag{S11}$$

When this condition is satisfied, the sign of $\dot{x}$ does not change in the range $0 < x < 1$, and this sign determines the long-term stable behavior. Choosing the representative point $x = 1/2$, we see that $\dot{x} = (1/4)\sigma_m(1/2)$, so we can use the sign of $\sigma_m(1/2) = s_m + (1/2)(\alpha_{mw} - \alpha_{wm})$: a positive sign means long-term stationary state will be at $x = 1$ while a negative sign means long-term stationary state will be at $x = 0$. Comparing this behavior to the non-interacting long-term behavior yields the conditions in the main text for maintenance, mirroring, and mimicry. Additionally, $\dot{x}$ can be identically zero if $\sigma_m(x) = 0$ for all $x$, which means $s_m + \alpha_{mw} = \alpha_{mw} + \alpha_{wm} = 0$; for $s_m \neq 0$, this constitutes masking.

### S6 Stochastic steady-state solution can be expressed in terms of interaction strengths

A form of steady-state solution $\rho^{\text{eq}}(x)$ for the Fokker-Planck-Kolmogorov equation in **Eq.** (B**3**) is

$$\rho^{\text{eq}}(x) = C\big[\exp\big(-U(x)\big)\big], \tag{S12}$$

where $C$ is a normalization constant, and we define the potential function $U(x) = \ln D(x) - \int_0^x dx'\, v(x')/D(x')$. For our interacting model, the integral can be carried out explicitly, yielding

$$U(x) = (1 - 2N\mu)\ln[x(1-x)] - 2N(s_m + \alpha_{mw})x + N(\alpha_{mw} + \alpha_{wm})x^2 - \ln(2N). \tag{S13}$$

Our solution describes a probability density function (PDF) for the steady-state mutant frequency for given population size, mutation rate, selection coefficient, and interaction terms. Knowledge of this $\rho^{\text{eq}}(x)$ can then be used to calculate summary values of the distribution – for example, the modal mutant population fraction or the mean $\bar{x} = \int_0^1 dx' x' \rho^{\text{eq}}(x')$. We show (**Figure S2**) that the game landscape is populated by probability density functions approximating the outcomes of the deterministic system, including bimodal distributions corresponding to bistable deterministic systems.

We observe (**Figure S4**) that for large populations under very weak interaction and at low mutation rates, the form of the solution is similar to the deterministic case (i.e. in the absence of noise), where the distributions approach delta functions at either $x = 0$ or $x = 1$. The values of $\alpha_{mw}$, $\alpha_{wm}$, and $\mu N$ alter the width of the peaks in the probability distributions.

### S7 Relationship between interaction coefficients for emulation in the presence of mutation and noise

We seek to classify maintenance, mirroring, masking, and mimicry in the presence of mutation and noise, in which the long-term behavior of the evolving population is described by the equilibrium probability distribution introduced in the previous section. We will summarise this distribution by its mode, i.e. its peak value, by which it is dominated in the limit as population size $N$ goes to infinity (we are in this limit when we use the Fokker-Planck-Kolmogorov description of Wright-Fisher dynamics). To begin to find the mode, we want to find all of the maxima and minima of the probability distribution, which can be found by setting the slope of the distribution equal to zero, or equivalently by setting to zero the derivative of the potential $U(x)$ defined above in **Eq.** (S13):

$$\begin{aligned} 0 = U'(x) &= (1-2N\mu)\left(\frac{1}{x}-\frac{1}{1-x}\right) - 2N(s_m+\alpha_{mw}) + 2N(\alpha_{mw}+\alpha_{wm})x \\ \Rightarrow 0 &= \left(\mu-\frac{1}{2N}\right) + \left[\alpha_{wm}+s_m-2\left(\mu-\frac{1}{2N}\right)\right]x + (-\alpha_{mw}-2\alpha_{wm}-s_m)x^2 + (\alpha_{mw}+\alpha_{wm})x^3 \end{aligned} \tag{S14}$$

Note that this is exactly the same as the cubic equation **Eq.** (S4) describing the long-term behavior of the RME, except mutation rate $\mu$ is replaced by $\tilde{\mu} = \mu - 1/2N$ (which we assume to be positive here, otherwise the probability distribution blows up at the boundaries). Thus the solution with no interactions is given by **Eq.** (S5) but with the replacement $\mu \rightarrow \tilde{\mu}$, which appears in the main text as the equation for the mode $x_{\text{mode}}(s_m)$ of the non-interacting stationary probability distribution with selection coefficient $s_m$ (the other zero of the quadratic equation is outside of the accessible range of $x$ between 0 and 1).

Observing the cubic equation **Eq.** (S14) and comparing it to its quadratic counterpart in the absence of interactions, we see that $x_{\text{mode}}(s')$ will be a zero of the cubic equation (meaning it will be either a minimum or maximum of the probability distribution) if $s' = \sigma_m^{s_m}(x_{\text{mode}}(s')) = s_m + \alpha_{mw} - (\alpha_{mw}+\alpha_{wm})x_{\text{mode}}(s')$ for some value $s'$. This relationship forms the basis of our emulation classification in the presence of mutation and noise; plots of these relationships for the different emulation categories are shown in **Figure S6**.

If, furthermore, this value $x_{\text{mode}}(s')$ is not just *a* maximum or minimum, but is in fact the unique maximum between 0 and 1. The interacting system truly emulates a non-interacting system with selection coefficient $s'$, since all non-interacting systems have a unique maximum between 0 and 1.

If we divide the cubic equation **Eq.** (S14) by the known solution factor $x - x_{\text{mode}}(s')$, this yields a quadratic equation for the two other zeros of the cubic besides $x_{\text{mode}}(s_m)$, with solutions:

$$x^{\pm} = \frac{1}{2}\left[1+\frac{\mu-1/2N}{\alpha_{mw}+\alpha_{wm}}\left(\frac{1}{1-x_{\text{mode}}}-\frac{1}{x_{\text{mode}}}\right) \pm \sqrt{\left[1+\frac{\mu-1/2N}{\alpha_{mw}+\alpha_{wm}}\left(\frac{1}{1-x_{\text{mode}}}-\frac{1}{x_{\text{mode}}}\right)\right]^2 + \frac{4(\mu-1/2N)}{\alpha_{mw}+\alpha_{wm}}\left(\frac{1}{x_{\text{mode}}}\right)}\right] \tag{S15}$$

To ensure $x_{\text{mode}}$ is the unique maximum between 0 and 1, we want to exclude areas of parameter space where these other potential maxima/minima $x^{\pm}$ are real and between 0 and 1. Upon inspection, this means we must enforce that $\alpha_{mw}+\alpha_{wm}$ is greater than or equal to the value that zeroes the expression inside the square root in $x^{\pm}$, which yields the constraint:

$$\alpha_{mw}+\alpha_{wm} \geq -(\mu-1/2N)\left[\frac{1}{\sqrt{x_{\text{mode}}}}+\frac{1}{\sqrt{1-x_{\text{mode}}}}\right]^2 \tag{S16}$$

### S8 Agreement of analytics with stochastic simulations across selection and interaction strengths

Where the outcomes of deterministic games are redundant in their contributing game interactions, the stochastic Fokker-Planck-Kolmogorov system produces a probability distribution dependent on N, $\mu$, s, $a_{mw}$, and $a_{wm}$. By simulating stochastic population dynamics, we can sample populations to create a probability distribution over the fractions of mutant and wild-type alleles.

The mean value of the simulation fractions is plotted in **Figure S2** for a specific $s_m(=0.05)$ with varying interaction coefficients, and the corresponding theoretical probability distributions in several cases are shown for comparison. It is worth noting that the "snowdrift" game from the lower left quadrant in traditional game space has two peaks in its stationary

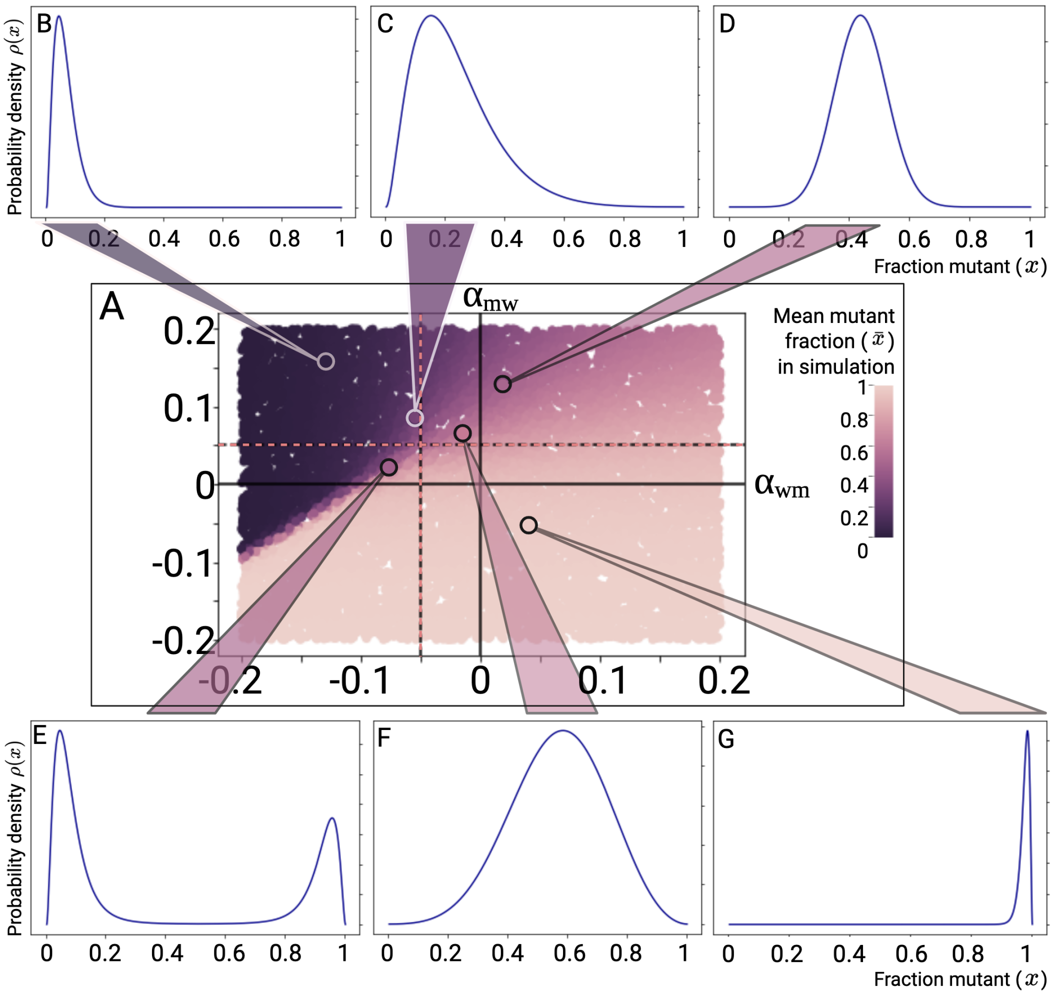


**Figure S2.** **Simulations and equivalent Fokker-Planck-Kolmogorov solutions for examples of a stochastic evolving and mutating population.** **(A)**The central plot displays the mean fraction of the mutant population ($\bar{x}$), plotted for 10,000 randomly distributed game parameters. Mean mutant fractions are plotted against randomly sampled interaction strengths, with mutant selection advantage $s_m = 0.05$, $N = 1000$, $\mu = 0.005$. The average (mean) proportion of mutant was plotted for each pair of interaction coefficients. Simulation results are colored by population fraction, with 100% wild-type, 0% mutant in dark purple, and 100% mutant, 0% wild-type population in cream. 10,000 random values of $\alpha_{mw}$, $\alpha_{wm}$ in the interval $[-0.2, 0, 2]$ were sampled to populate the phase plot. Six distributed examples (**B-G**) of the analytical Fokker-Planck-Kolmogorov equilibrium solution for $\rho(x)$ against $x$ are shown and their approximate position in the simulation space is highlighted. Distributions from the upper left and lower right (e.g., **B**, **G**) have distributions strongly peaked near 0% and 100% mutant fractions, respectively. The upper right quadrant (e.g., **D**, **F**) shows stable co-existence, with solutions and simulations in the bottom left quadrant (e.g., **E**) representing probability distributions with two peaks, one near 100% wild-type and one near 0% wild-type.

Fokker-Planck-Kolmogorov solution, and the height of these peaks becomes more uneven with distance away from the line $\alpha_{wm} = \alpha_{mw} - 2s_m$. The width of the peaks in all sections increases with mutation rate $\mu$.

In the Wright-Fisher model with mutation, a population under strong selection will move to the peak of the landscape in a single-peaked landscape, whereas in a flat landscape, it fluctuates stochastically around equal proportions of all genotypes. Without interactions, the results of the Wright-Fisher model depend only on the intrinsic (genetic) fitness landscape – in particular, whether the landscape is neutral or peaked – and the mutation rate. To validate the predicted result of interactions in evolutionary simulation models we added randomly distributed interaction strengths to a Wright-Fisher model of both single-peaked ($s_m \neq 0$) and flat ($s_m = 0$) landscapes.

In **Figure** S3 and **Figure** S5 we examine the interacting population's probability density function at stationarity. In **Figure** S3**A** we plot the modal values of the analytical solution over varying $\alpha_{mw}$ and $\alpha_{wm}$ for a given selection and mutation space. To validate these distributions we sampled the frequency spectrum of the population in simulations at long time points (between 100 and 10,000 generations). We compared this frequency distribution to our analytical probability density equilibrium solution for the Fokker-Planck-Kolmogorov equation with interactions.

The equilibrium distribution of a system with known monoculture fitnesses and no game interactions is well-defined and

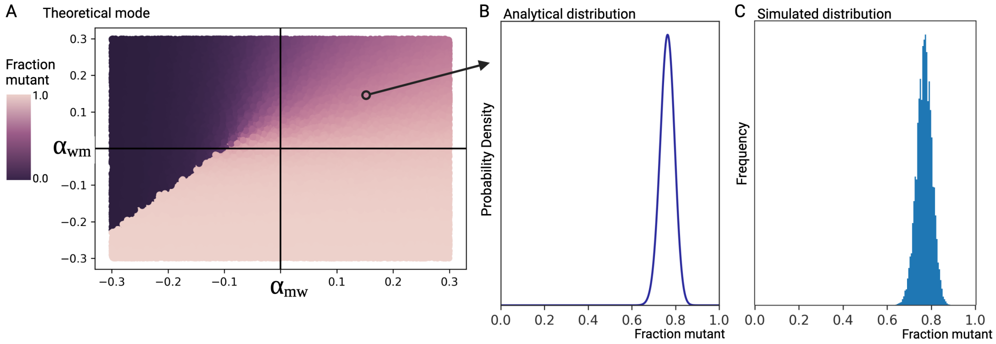


**Figure S3. Fokker-Planck (FP) distribution compared to stochastic simulation results. (A)** The modal value of the analytical FP distribution is plotted for 100,000 randomly generated pairs of interaction terms. Modal values of 100% wild-type are dark purple and 100% mutant fraction are colored cream. **(B)** We highlight the shape of the analytical probability density distribution in a specific Fokker-Planck solution ($\alpha_{wm} = 0.16,\ \alpha_{mw} = 0.14$, $s_m = 0.1$) and compare theoretical results in plot **(B)** to simulation. **(C)** Distribution of sampled fractions between 200 to 500 generations in 500 independent simulations is shown. These simulations had a selection coefficient of $s_m = 0.1$, representing a fitter mutant population. The histogram is a sampled distribution of the population fraction, with each sample measured at $t > 200$ generations from 500 simulations under these conditions. Further examples are shown in **Figure S5**.

understood in population genetics. As a result, the equilibrium distributions/evolutionary outcomes measured in experiments are often assumed to result from entirely intrinsic fitness differences. This assumption does not account for potential interactions between populations. As seen in **Figure S2**, the survival of the fittest (under which the 'fittest' genotype prevails) in the presence of interactions becomes frequency-dependent and can result in multiple populations co-occurring.

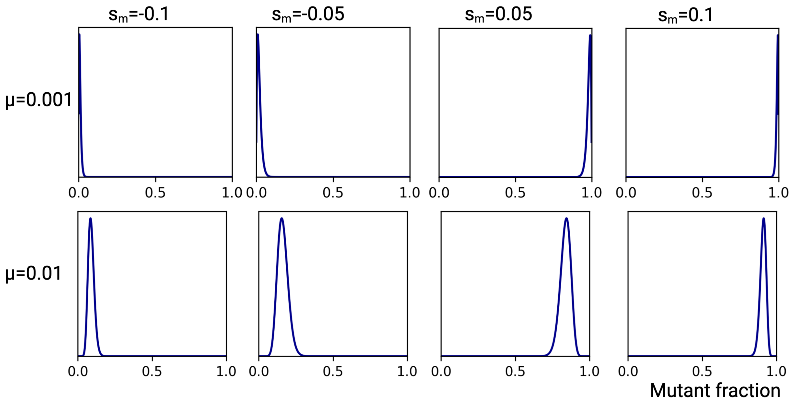


**Figure S4. Fokker-Planck analytical probability density distribution for minimal interaction strengths agrees with baseline expectations.** Each example represents the analytical distribution for a given population size (N=1000) and given small interaction strengths ($\alpha_{wm} = \alpha_{mw} = 0.0001$) for a range of mutant selection coefficients $s_m$, and mutation rates $\mu$.

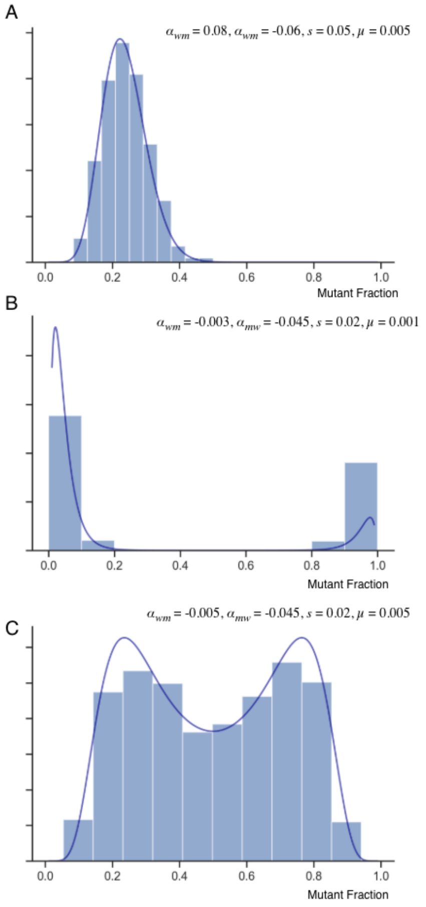


**Figure S5. Fokker-Planck-Kolmogorov solution for probability density holds under simulations of a range of interaction parameters.** Each example represents the analytical FPK solution (dark blue line, overlay) and a histogram of sampled mutant fraction at generation 1500 for 2000 independent simulations. **(A)** $s_m = 0.05$, $\alpha_{wm} = 0.08$, $\alpha_{mw} = -0.06$, $\mu = 0.005$, $N = 1000$. **(B)** $s_m = 0.02$, $\alpha_{wm} = -0.003$, $\alpha_{mw} = -0.045$, $\mu = 0.001$, $N = 1000$. **(C)** $s_m = -0.05$, $\alpha_{wm} = -0.045$, $\alpha_{wm} = -0.005$, $\mu = 0.005$, $N = 1000$.

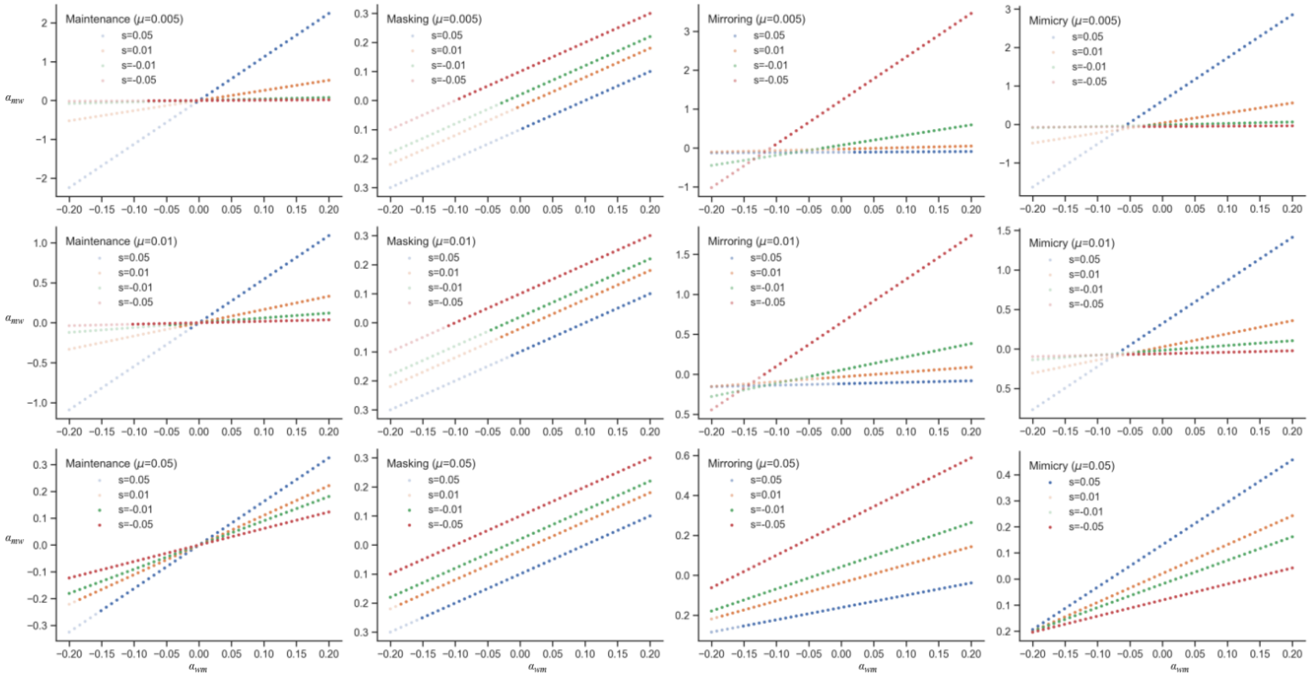


**Figure S6. Relationship between interaction coefficients.** The relationships between $\alpha_{wm}$ and $\alpha_{mw}$ as defined analytically in the main text. Functions for maintenance, masking, mirroring, and mimicry are plotted for three fixed mutation rates and varying intrinsic selection coefficients. Transparency is used to distinguish parameter combinations that meet the condition of a solution with a unique maximum (solid) or multiple (transparent).

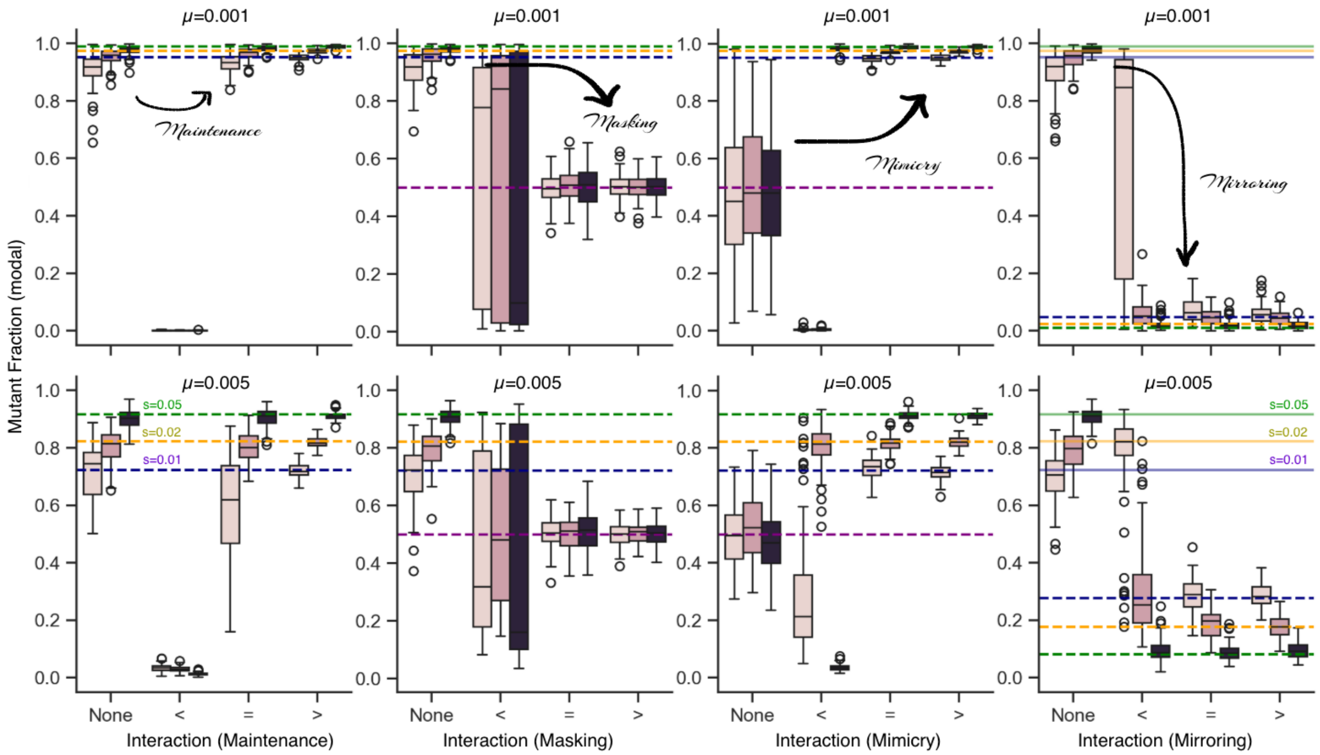


**Figure S7. Masking, mimicry, maintenance, and mirroring predictions hold under parameter alteration.** Further simulations of maintenance, masking, mirroring, and mimicry for different mutation rates and different intrinsic selection coefficients. Simulations are ordered left to right by no interactions (None, intrinsic selection only), insufficient interactions ($<$), minimal interaction strength for the effect to manifest and meet the conditions for unique maxima ($=$), and increased strength interactions ($>$). Each boxplot represents 100 simulations for a given set of selection, interaction, and mutation strengths.

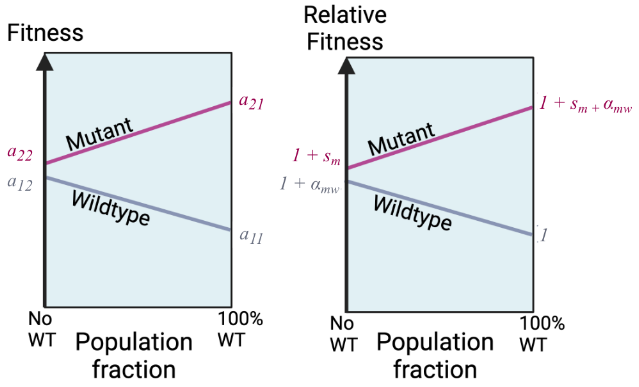


**Figure S8.** **Illustration of how the fitness-frequency plot**. This plot, constructed similarly to an invasion diagram, can be rewritten in terms of intrinsic and interaction terms. **(Left)** The payoff matrix (similar to the invasion diagram) consists of the values of the four intersections of the frequency-fitness lines with the axes. While the 100% WT or mutant fractions can be directly measured, the infinitesimally small $\rightarrow$ 0% growth rate of the minority population is impossible to measure directly and must be inferred using regression and assumptions (of linearity or otherwise) from the other values. **(Right)** The equivalence of the intersections in terms of the modified payoff matrix is shown where the growth rates are normalized to the wild type. In the traditional game plot, the differences between the intersections are plotted, the two resultant values are functions of 3 variables and thus form a many-to-one mapping.

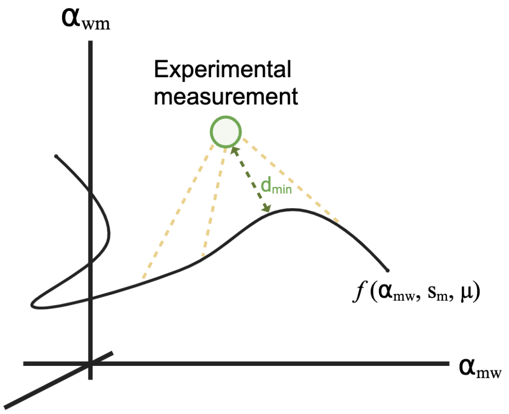


**Figure S9. Illustration of Euclidean distance of the experimental data points from a theoretical regime.** Distance $d_{\text{min}}$ is calculated via numerical optimization (Python) to find the minimum distance between the point representing the experimental result $(\alpha_{wm}, \alpha_{mw}, s, \mu)$ and all points constrained to the specific manifolds $(\alpha_{wm} = f(\alpha_{mw}, s, \mu))$ that represent the theoretical conditions of masking, maintenance, and mirroring.

### Supplemental Tables

**Table S1.** Additional details of the published experiments used to derive payoff matrices

| Author | Year | System | Variable | Payoff matrices | Data Type |
|---|---|---|---|---|---|
| Vulic and Kolter [4] | 2001 | Bacteria | *Escheria coli* WT and GASP mutant | 2 | Growth-Frequency Plot |
| Li et al. [5] | 2015 | Bacteria | *Curvibacter* sp. (AEP1.3) and *Duganella* sp. (C1.2) | 1 | Growth-Frequency Plot |
| Kaznatcheev et al. [1] | 2019 | Cancer | NSCLC Cell Lines Drug+Fibroblasts | 4 | Payoff matrix |
| Adamowicz et al. [6] | 2020 | Bacteria | *Escheria coli*, *Salmonella enterica* *mdoG* and *mdoH* gene mutants | 3 | Growth-Frequency Plot |
| Cai et al. [7] | 2020 | In-silico | Metabolic simulation | 1 | Payoff |
| Faroukkian et al. [8] | 2020 | Cancer | NSCLC Cell lines Mutation, Drug | 5 | Payoff |
| Maltas et al. [9] | 2024 | Cancer | NSCLC Cell Line Mutation, Drug | 4 | Payoff |

**Table S2.** Payoff values and equivalent selection and interaction terms extracted from papers

| | Paper | $a_{11}$ | $a_{12}$ | $a_{21}$ | $a_{22}$ | $s$ | $\alpha_{wm}$ | $\alpha_{mw}$ |
|---|---|---|---|---|---|---|---|---|
| 0 | Kaznatcheev2019 | 2.6 | 3.5 | 3.1 | 3.0 | 0.1538 | 0.3462 | 0.0385 |
| 1 | Kaznatcheev2019 | 2.5 | 2.4 | 4.0 | 2.7 | 0.08 | -0.04 | 0.52 |
| 2 | Kaznatcheev2019 | 0.5 | -0.4 | 3.8 | 2.4 | 3.8 | -1.8 | 2.8 |
| 3 | Kaznatcheev2019 | 2.3 | 4.3 | -1.3 | -1.0 | -1.4348 | 0.8696 | -0.1304 |
| 4 | Maltas2024 | 1.0 | 0.97 | 0.97 | 0.84 | -0.16 | -0.03 | 0.13 |
| 5 | Maltas2024 | 1.03 | 0.99 | 1.01 | 0.93 | -0.0971 | -0.0388 | 0.0777 |
| 6 | Maltas2024 | 1.0 | 1.02 | 0.95 | 0.88 | -0.12 | 0.02 | 0.07 |
| 7 | Maltas2024 | 1.01 | 1.01 | 0.98 | 0.9 | -0.1089 | 0.0 | 0.0792 |
| 8 | Farrokhian2022 | 0.9963 | 1.004 | 0.975 | 0.7517 | -0.2455 | 0.0077 | 0.2241 |
| 9 | Farrokhian2022 | 0.47 | 0.42 | 0.83 | 0.7 | 0.4894 | -0.1064 | 0.2766 |
| 10 | Farrokhian2022 | 0.377 | -0.01 | 0.86 | 0.714 | 0.8939 | -1.01 | 0.3873 |
| 11 | Farrokhian2022 | 0.37 | -0.43 | 0.93 | 0.728 | 0.9676 | -2.1622 | 0.5459 |
| 12 | Farrokhian2022 | 0.33 | -0.59 | 1.12 | 0.789 | 1.3909 | -2.7879 | 1.003 |
| 13 | VulicKolter2001 | 1.0 | 0.2 | 1.1 | 0.5 | -0.5 | -0.8 | 0.6 |
| 14 | VulicKolter2001 | 0.5 | 1.1 | 0.2 | 1.0 | 1.0 | 1.2 | -1.6 |
| 15 | CaiChan2020 | 1.18 | 1.09 | 1.42 | 1.34 | 0.1356 | -0.0763 | 0.0678 |
| 17 | Li2015 | 0.1774 | 0.1284 | 0.13895 | 0.08 | -0.549 | -0.2762 | 0.3323 |
| 18 | Adamowicz2020 | 0.89 | 0.37 | 0.15 | 0.61 | -0.3146 | -0.5843 | -0.5169 |
| 19 | Adamowicz2020 | 0.45 | 0.4 | 0.43 | 0.6 | 0.3333 | -0.1111 | -0.3778 |
| 20 | Adamowicz2020 | 0.64 | 0.39 | 0.41 | 0.29 | -0.5469 | -0.3906 | 0.1875 |
